# Brain-like neural dynamics for behavioral control develop through reinforcement learning

**DOI:** 10.1101/2024.10.04.616712

**Authors:** Olivier Codol, Nanda H. Krishna, Guillaume Lajoie, Matthew G. Perich

**Author notes:** Co-senior authors (Guillaume Lajoie, Matthew G. Perich).

## Abstract

During development, neural circuits are shaped continuously as we learn to control our bodies. The ultimate goal of this process is to produce neural dynamics that enable the rich repertoire of behaviors we perform. What begins as a series of “babbles” coalesces into skilled motor output as the brain rapidly learns to control the body. However, the nature of the teaching signal underlying this normative learning process remains elusive. Here, we test two well-established and biologically plausible theories—supervised learning (SL) and reinforcement learning (RL)—that could explain how neural circuits develop the capacity for skilled movements. We trained recurrent neural networks to control a biomechanical model of a primate arm using either SL or RL and compared the resulting neural dynamics to populations of neurons recorded from the motor cortex of monkeys performing the same movements. Intriguingly, only RL-trained networks produced neural activity that matched their biological counterparts in terms of both the geometry and dynamics of population activity. We show that this similarity with biological brains depends critically on matching biomechanical properties of the limb. Dynamical analysis on network activity revealed that our RL-trained networks operate at the “edge of chaos”, a dynamical regime known for its computational richness, greater memory capacity, and robust plasticity properties. We then demonstrated that monkeys and RL-trained networks, but not SL-trained networks, show a strikingly similar capacity for robust short-term behavioral adaptation to a movement perturbation, indicating a fundamental and general commonality in the neural control policy. Together, our results support the hypothesis that neural dynamics for behavioral control emerge through a process akin to reinforcement learning. The resulting neural circuits offer numerous advantages for adaptable behavioral control over simpler and more efficient learning rules and expand our understanding of how developmental processes shape neural dynamics.

## Introduction

During development, neural circuits are shaped continuously as we learn to control our bodies. In the first days and weeks of an infant’s life, movements appear to a nearby spectator as “babbling”; a disorganized series of twitches and tumbles without purpose or intent. Yet, the brain quickly leverages the sensorimotor information gathered through that period to build an efficient and versatile control policy for the biomechanics of each individual’s body. This control policy ultimately shapes the neural dynamics that underlie behavior in regions of the primate brain such as the motor cortex^1–4^. This normative learning process requires optimization of neural circuits using a teaching signal that encapsulates the success of a movement relative to its goal^5–9^ (e.g., to grab food or beckon a parent). While the nature of that signal remains elusive, there are two well-established and biologically plausible theories (Fig. 1). The brain might learn directly from the errors made during an attempted action^10,11^: is the arm reaching towards a piece of fruit, or must the output be adjusted? Additionally, the brain might learn by evaluating the success of the actions *post hoc*^6,12–15^: did the arm successfully grab the fruit, or must I adjust my planned action on the next attempt? Here, we ask what is the learning scheme that best captures the dynamics and representations found in the brain.

**Fig. 1.**
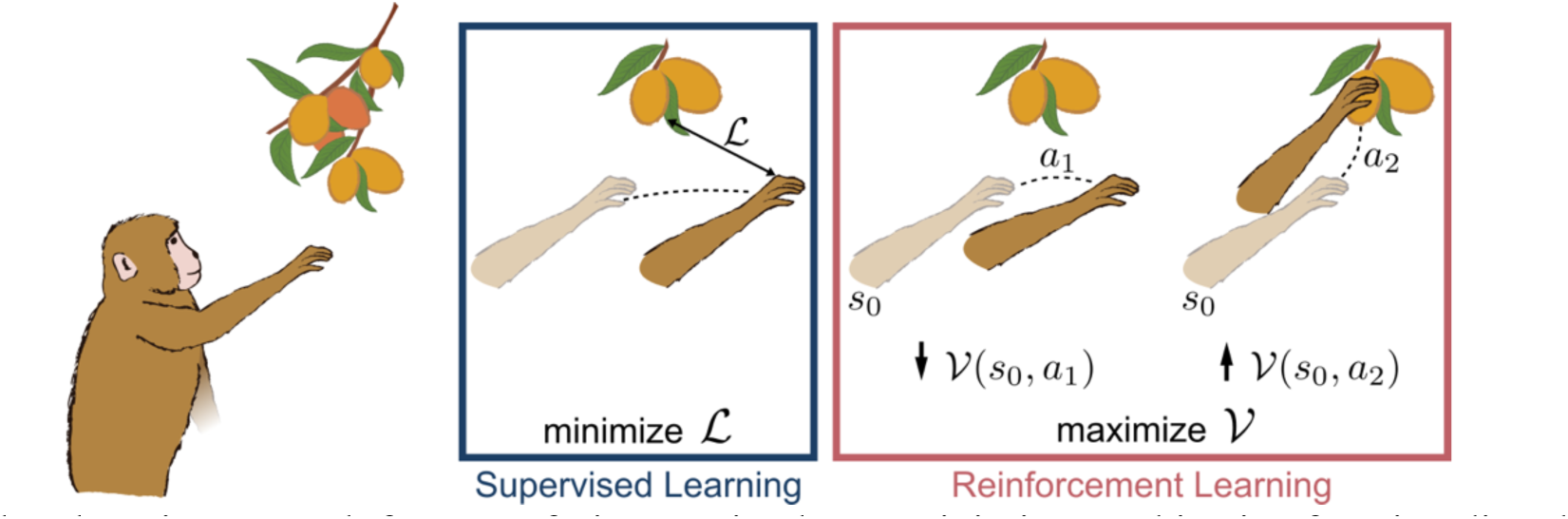
When learning to reach for some fruit, an animal may minimize an objective function directly (Supervised Learning, left), or learn the value of a movement and maximize the surrogate value function instead (Reinforcement Learning, right).

We formalize the first theory as a supervised learning (SL) process, where the teaching signal is the direct gradient of an error function, or loss to be minimized; in this case the error between intended and performed actions^11^. Updating synaptic weights of a neural circuit proportionally to this gradient is the backbone of SL and is widely used for deep learning. This framework is computationally compelling because plasticity is driven by a direct measure of the goal, giving efficient and rapid convergence to an optimal solution for the control policy. However, SL requires an explicit knowledge of the relationship between the loss function and the synaptic weights undergoing plastic changes which is difficult to ascertain in real neural circuits (though numerous implementations have been proposed^10^).

The second theory can be formalized through a reinforcement learning (RL) process. In contrast to SL, where errors are accurately and continuously defined, RL proposes that the central nervous system estimates the value (e.g., success) of an action after it has been performed and adjusts the synaptic weights to maximize reward^15–17^. This is achieved by defining a surrogate reward function that approximates the true loss function, allowing gradient-based optimization on the intermediate reward function rather than the error directly^16^. Consequently, RL is in practice a noisier process than SL, leading to slower and less robust convergence in general.

However, the RL framework is appealing in the context of learning in biological organisms. RL conceptually encapsulates our intuition about development where initial movements are made seemingly at random and without intention^18–21^. Successful actions like grabbing a mother’s hand could be selectively reinforced without an explicit goal. RL is also computationally desirable for biological implementations, as it does not require explicit knowledge of the loss gradient, overcoming the major biological limitation of SL^10,11,16^. Indeed, there is intriguing evidence supporting RL-like processes in the brain. Many neural regions exhibit online encoding of reward, including the striatum^22–24^, the prefrontal cortex^25–27^, and even the motor cortex^27–31^. Some regions, such as prefrontal cortex, exhibit both context-dependent^32–34^ and intrinsic (motivational)^35,36^ modulation with reward. This flexibility is critical to develop viable control policies in biological agents, where information from an ever-changing environment must be efficiently integrated into online control of movement to produce adequate behaviour^37–39^. Finally, maximizing reward is a transferable learning framework across environments and species. For instance, “securing food” is a context-agnostic reward signal, whether it be by reaching up for an apple or down for a strawberry on a bush. Therefore, RL provides a smooth optimization path for learning at both developmental and evolutionary timescales^40,41^.

Here, we directly compare these two theories as models of the development of behavioral control. Given the conceptual and computational advantages outlined above, we hypothesized that RL-like processes would better explain the properties of neural population activity in biological brains. We tested this hypothesis by comparing neural population recordings from the motor cortex of macaque monkeys performing reaching movements to the activity of *in silico* neural network models trained to produce the same behaviors using either RL or SL^42,43^. We assessed similarity through both the geometrical and dynamical properties of the population activity to capture multiple facets of the underlying neural computations^2,44–46^. We show that despite achieving similar performance in behavioral control, the neural activity of these models is easily distinguishable and neural recordings of monkeys better align with RL activity. Critically, we show that these differences between RL and SL models are dependent on the biomechanical properties of the effector being controlled^4,43^, and that RL-trained networks operate at the “edge of chaos”, a dynamical regime at the transition zone between stability and instability that has long been considered advantageous for biological neural circuits^47,48^. Finally, we altered the environmental dynamics following learning and demonstrate that this led our RL-trained models to undergo reorganization of neural activity that mirrors biological reorganization^49^, while our SL-trained models failed to do so. Together, these findings indicate that the neural control solutions implemented in the central nervous system are likely shaped in large part by reinforcement learning signals. The resulting circuit offers numerous advantages for robust behavioral control over simpler and more efficient learning rules and has important implications for our understanding of how developmental processes shape behaviorally relevant neural dynamics^18,40,41^.

## Results

### Neural networks learn to produce successful movements through both RL and SL

We designed a simulated environment to test theories of teaching signals for *de novo* acquisition of motor skills. Using the MotorNet toolbox^43^, we instantiated recurrent neural network models that were directly linked to two-link biomechanical models of the primate arm driven by six muscle actuators. We trained the models to perform reaching movements using one of two learning rules: direct gradient descent on output error during the movement (SL; Fig. 2a), or reinforcement learning implemented with Proximal Policy Optimization^50^ (RL; Fig. 2b). Except for the learning rule, all models were comprised of identical features, including RNN architecture, musculoskeletal arm, and task-relevant inputs (Fig. 2c). Critically, the reward function provided to SL models was same loss function as employed in RL models without the negative sign to accommodate for gradient descent rather than gradient ascent. This ensured the normative quantification of the task goal being optimized remained identical across models. Therefore, any differences in neural representation or dynamics were exclusively driven by the differences in each model’s learning rule. We trained the models *de novo*, from random initializations, to perform a set of random point-to-point reaches. Both model families learned to effectively control the arm, achieving high performance (n=24 seeds per group, seed-matched across group; Fig. 2d-e). Neural recordings to model comparisons were done at epochs with matching evaluation performance to rule out training bias.

**Fig. 2:**
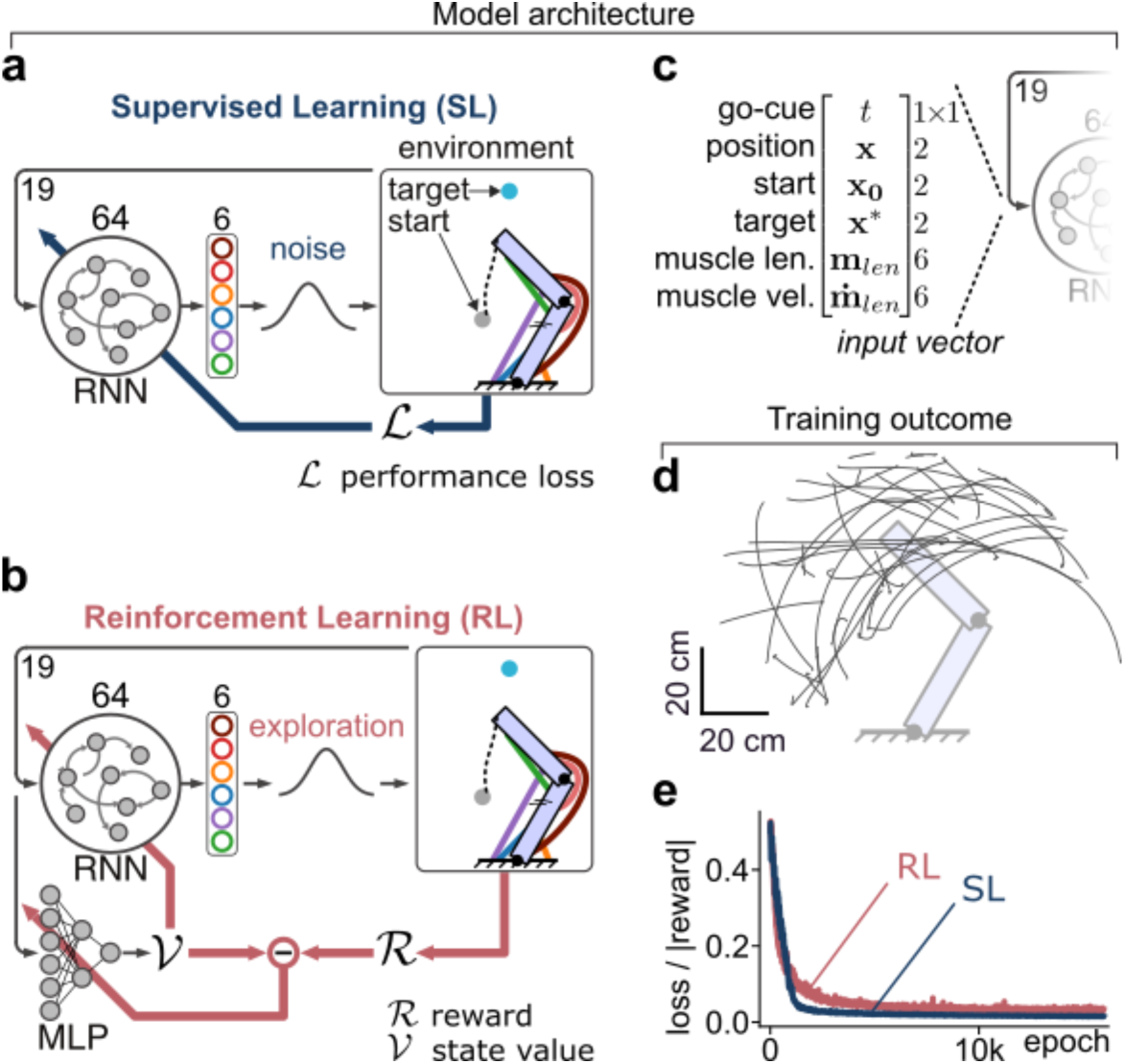
Neural networks trained with RL or SL achieved high performance in controlling the effector. **a**, **b**. Neural networks consisted of a recurrent neural network (RNN) containing 64 Gated Recurrent Units projecting to a fully connected output layer. A Gaussian noise process was added to the output of all models, which served to drive exploration for RL models (panel b). In RL models a separate multi-layer-perceptron (MLP) acted as a critic to learn a value function (panel b). **c**. Neural networks received task-relevant inputs about the goal and instantaneous state of the effector. **d**. Example post-training point to point reaching trajectories along the full joint space. **e**. All neural networks learned to minimize the loss (SL, blue) or equivalently maximize reward (RL, red). Note that the reward function is strictly equivalent to the negative of the supervised loss (see Methods for details).

### RL-trained networks show high geometric similarity to population activity in monkey motor cortex

We sought to compare the activity of the RL- and SL-trained networks to determine if one group better matched the activity observed in biological brains. We analyzed recordings of neural populations in the primary motor (M1) and premotor (PMd) cortex (Fig. 3a) of monkeys (male, *macaca mulatta*) performing a point-to-point, center-out planar reaching task. In brief, the monkeys reached to one of 8 targets positioned in a circle 8 cm from the center. We trained 24 seed-matched neural networks to perform random reaches using RL or SL (Fig. 2d), before freezing the weights and instructing the networks to perform the same task as the monkeys (Fig. 3b). We saw that all three networks—biological, RL, and SL—performed the task adeptly and successfully acquired the targets (Fig. 3c). We used Principal Component Analysis (PCA) to identify the leading 20 components of the neural activity in each dataset (Fig. 3d-e). Intriguingly, we saw similar target organization between the biological and RL-trained, but not SL-trained, networks. We quantified this geometric similarity using Canonical Correlation Analysis (CCA) on these leading 20 principal components of each dataset (Fig. 3e-f). Remarkably, RL models showed higher canonical correlation values than SL models when compared against monkey neural recordings (Fig. 3f-g; rank-sum test, monkey C: U=5.92, U=-4.53, U=-5.87 for RL-SL, RL-neural, and SL-neural comparisons, respectively; monkey M: U=5.92, U=-5.47, U=-5.87; p<0.001 for all comparisons), nearly approaching the upper bound set by across-monkey comparisons. This result clearly shows that the geometrical properties of neural activity in the RL models are more similar to neural activity in biological neural networks during the same task.

**Fig. 3:**
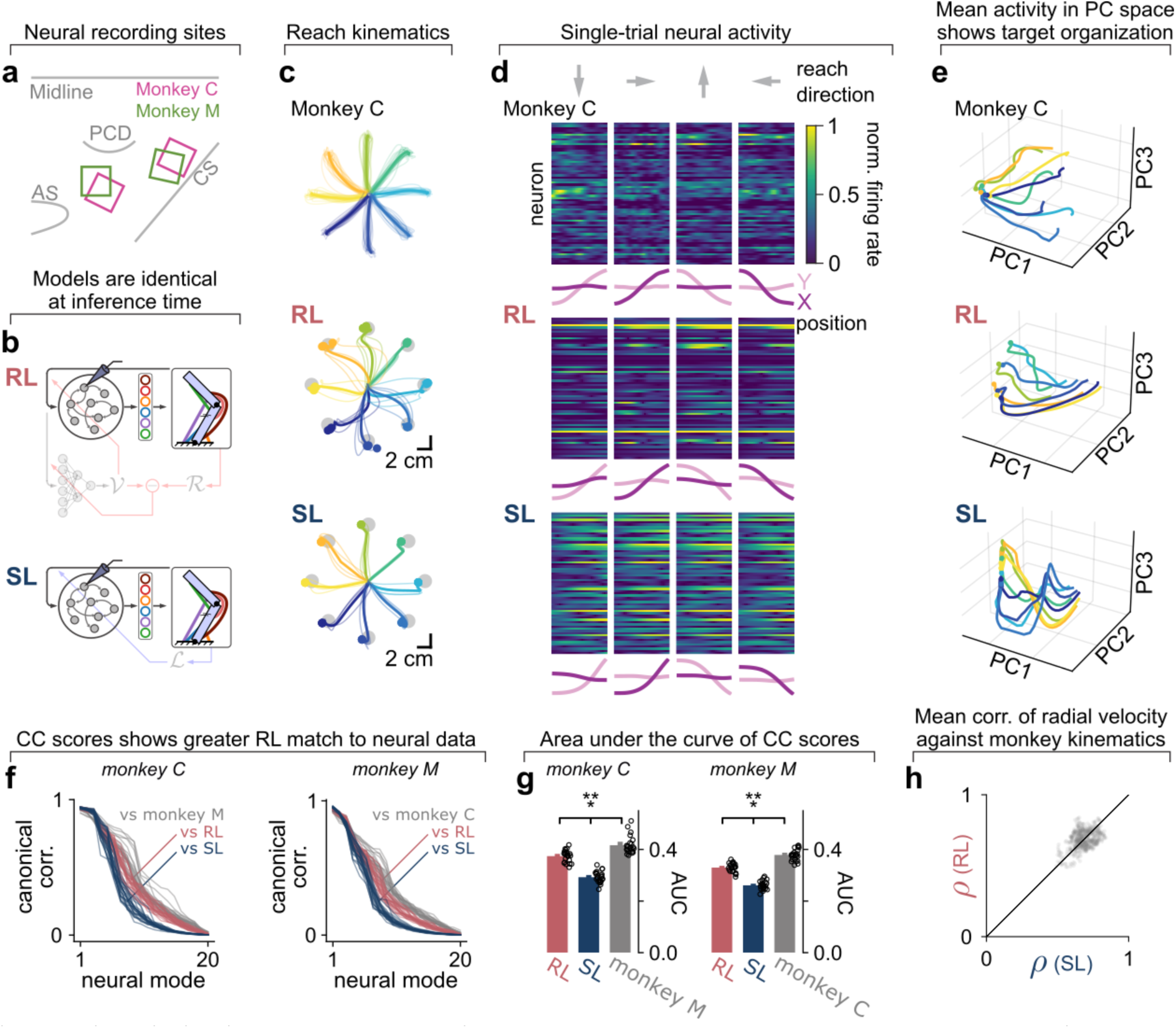
High similarity between RL-trained neural networks and monkey neural recordings. **a**. We recorded neural activity from primary motor cortex and premotor cortex of two males *macaca mulatta* monkeys using Utah arrays. **b**. The weights of our neural network models were frozen before being instructed to perform the same center-out reaching task as the monkeys. **c**. Example kinematics for one monkey and two models performing center-out reaches. The thin and thick lines indicate individual and mean reaches, respectively. **d**. Example normalized neural activity for one monkey and two neural network models for a trial in one of four cardinal reaching directions. Pink lines indicate kinematics during that trial. **e**. Neural trajectories of the leading 3 Principal Components after within-condition trial averaging. Note that averaging was performed after PCA. The dot at the end of each trajectory line indicates the final neural state. **f**. Canonical Correlation curves for each model against neural recordings. Each line represents a model seed (n=24) being compared to the same recording session. The grey lines represent comparison to different recording sessions from the other monkey (n=23). **g**. Area under curve (AUC) of the CC curves from panel f. Bar heights and error bars indicate the mean and 95% confidence intervals, respectively. **h**. Pearson correlation coefficients of condition-averaged radial velocity between monkey and neural network kinematics, plotted against unity based on neural network seed matching.

To verify that the similarity between RL-trained and biological networks was not due to closer behavioural match between model output and monkey kinematics, we compared the mean correlation value of radial velocities between of model and monkey reaches and did not find any difference (Fig. 3h; signed-rank test, monkey C: T(6)=5035, p=0.71; Monkey M: T(18)= 45920; p=0.75). The similarity of RL-trained networks to monkeys was robust to changes in hyperparameters of the neural network models including network size, activity regularization, exploration sampling rate, or batch size and was not dependent on our assumed dimensionality found through PCA (Fig. S1-S6). This was also robust to a change in synaptic learning rules from backpropagation through time to eligibility propagation^51^, demonstrating that our results specifically pertain to the nature of the teaching signal (Fig. S7).

The SL models benefit from having access to the exact loss^11^ (Fig. S8a-b), while RL models rely on an initially biased estimate of the value function that converges over time to an unbiased estimate^52^. To rule out the possibility that this difference could drive our main results, we took the critic networks post-training in our RL models, and re-trained new RL models using these pre-trained critics with their weights frozen (Fig. S8c-e), ensuring that they had early access to an accurate value estimator. Nevertheless, our canonical correlation results remained similar (Fig. 8f, rank-sum test, monkey C: U=4.82, U=-5.87, U=-5.87 for RL-SL, RL-neural, and SL-neural comparisons, respectively; monkey M: U=5.46, U=-5.72, U=-5.87; p<0.001 for all comparisons). This confirmed that our observations are not a consequence of the early availability of an accurate loss estimator during training and instead reflect a fundamental outcome of developing neural networks with RL. Interestingly, we note that providing a pre-trained critic to the RL models achieved straighter and more consistent reaches than those displayed by RL models which had to learn the critic concomitantly (Fig. S8d).

### RL-trained networks are dynamically similar to monkey motor cortex

Neural population activity is thought to arise from dynamical rules that determine how neural state trajectories smoothly evolve over time. These dynamical rules reflect the computations performed by the network. There thus remains the possibility that RL and SL models display different geometrical features in their neural activity but perform the same underlying computation to solve the control problem. We applied dynamical systems theory next to directly test the similarity of the underlying neural computations between the neural network models and monkey motor cortex. We leveraged Dynamical Similarity Analysis^53^ (DSA), which employs Koopman theory to reformulate a non-linear dynamical system as a higher-dimensional linear system via a mapping function *g* (Fig. 4a). This transformation preserves the dynamical motifs of each system, allowing us to use tractable, linear metrics such as Procrustes dissimilarity to quantify the difference between each system’s dynamical rules (Fig. 4a).

**Fig. 4:**
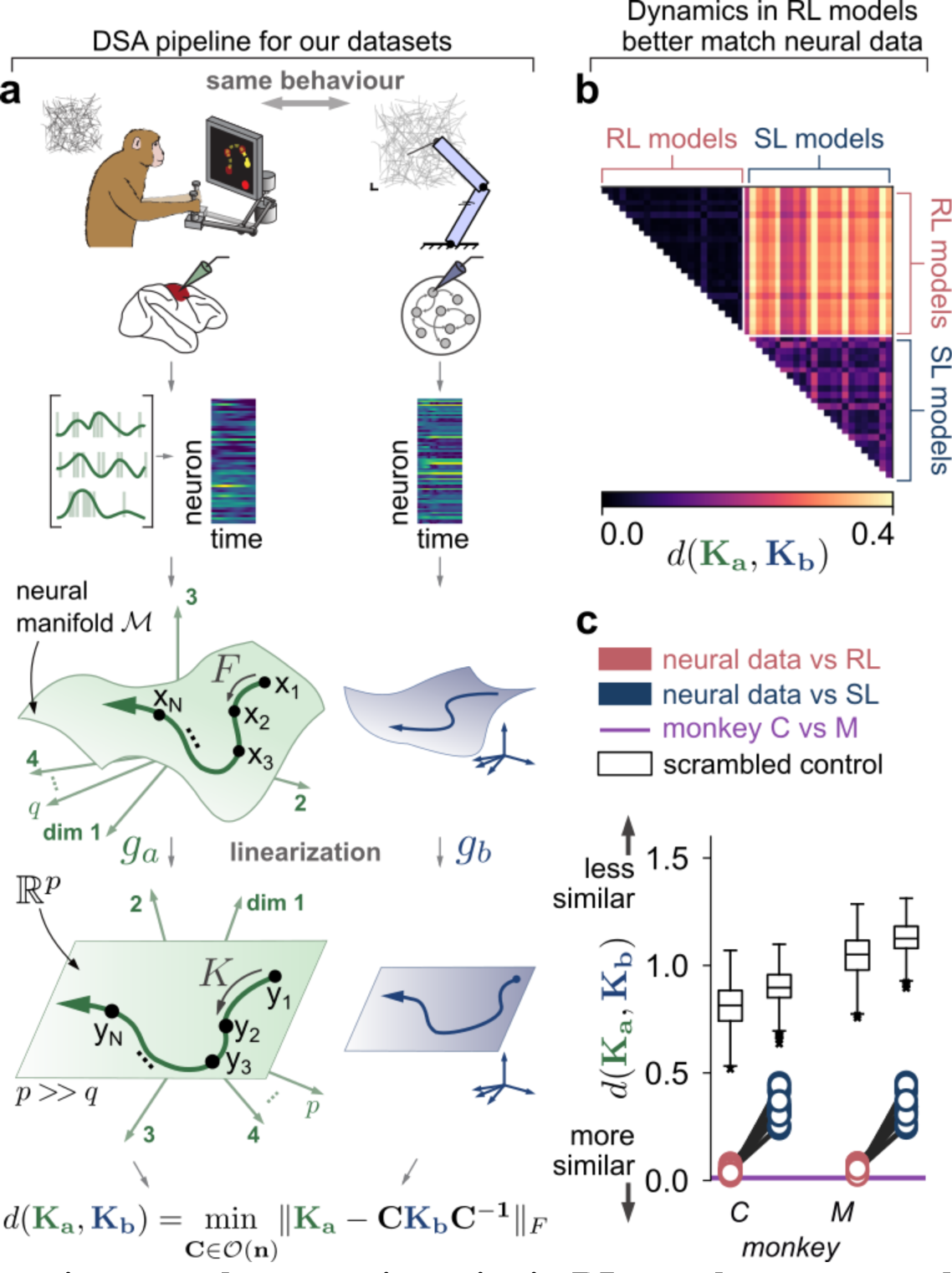
Dynamics governing neural state trajectories in RL are closer to neural recordings in motor cortices. **a**. Neural activity from motor cortices and models during a random target task were compared. Neural firing rates were linearized using Dynamical Similarity Analysis and their dissimilarity was quantified using a Procrustes metric over vector fields (see Methods and Ref. ^53^ for details). **b**. Dissimilarity scores between neural networks models (n=24 per group). Each row and column of the dissimilarity matrix represents a neural network seed. RL and SL seeds were ordered identically. **c**. Dissimilarity scores between neural network models and neural recordings from monkeys performing the random target task. The solid horizontal purple line represents dissimilarity between the two monkey recordings. The black box plots indicate the dissimilarity values for the same comparisons with the time dimension scrambled. The box and the whiskers indicate the quartiles and 1.5 times the interquartile range, respectively.

DSA requires a thorough sampling of the neural activity space to yield reliable estimates of the neural dynamics manifold^53^. While the center out task used in the previous analysis guarantees clear and interpretable geometric organization in the population activity, the task has the drawback that the networks explore only a limited portion of the possible activity state space. We therefore recorded motor cortical data from monkeys performing a more complex random point-to-point planar reaching task (Fig. 4a), where the start and end position of the reach are stochastically drawn from a distribution (see Methods). We then instructed our 24 seed-matched RL- and SL-trained neural networks to perform the exact same movements as the monkeys. We quantified the dynamical similarity of the computations performed by the biological and artificial networks using Procrustes dissimilarity in the aligned manifold found by DSA, where lower values indicate more similar dynamics. Interestingly, we found clear differences in dynamics between networks trained with RL versus those trained with SL (Fig. 4b; rank-sum test, U=-28.90, p<0.001), despite the fact all networks produced the same behavioral output. This indicates that the learning rule fundamentally shapes the computations performed by the network to produce that behavior.

Remarkably, when we compared these networks to monkey motor cortex, we found that RL produced significantly more brain-like dynamics than SL (Fig. 4c; sign-rank test, T=0, p<0.001 for both monkeys). To confirm the generalization of these findings, we repeated the analyses using data from the center-out task and saw the same qualitative result (Fig. S9; U=-25.85, p<0.001 for model-model comparisons; T=0, p<0.001 for both monkeys in model-recording comparisons). To contextualize the numbers, we compared to lower and upper bounds set by the neural data: the dynamical dissimilarity across different monkeys and the dynamical dissimilarity expected by chance, respectively. Note that although there remain some differences between RL networks and monkey motor cortex, dynamical alignment is much harder to achieve than geometrical alignment. Indeed, only one dynamical motif will minimize dynamical similarity, while there are a near-infinite number of solutions which could produce geometrically identical solutions, making it more striking that our RL-trained networks even partially align with the biological data.

### Brain-like neural dynamics stem from biomechanically realistic effectors

During *de novo* learning, neural circuits are optimized for the environment and behaviors they need to perform but must do so within the constraints of the effector with which they interact with the environment. In the case of primate reaching, this effector is the arm, which co-evolved with the brain over many generations^4,40^. We hypothesized that the specific biomechanical complexities of the effector should thus impact the similarity between the RL and biological networks. To test this hypothesis, we swapped the effector from a two-joint arm actuated by non-linear Hill-type muscles^54^ to a point-mass actuated by four linear muscles^43^ (Fig. 5a). This led the CC scores in RL-trained models to largely drop when compared to neural recordings of monkeys performing a center-out reaching task (Fig. 5b), approaching the CC scores of SL-trained models (Fig. 5c; rank-sum test, monkey C: U=1.51, p=0.13; monkey M: U=1.13, p=0.25).

**Fig. 5:**
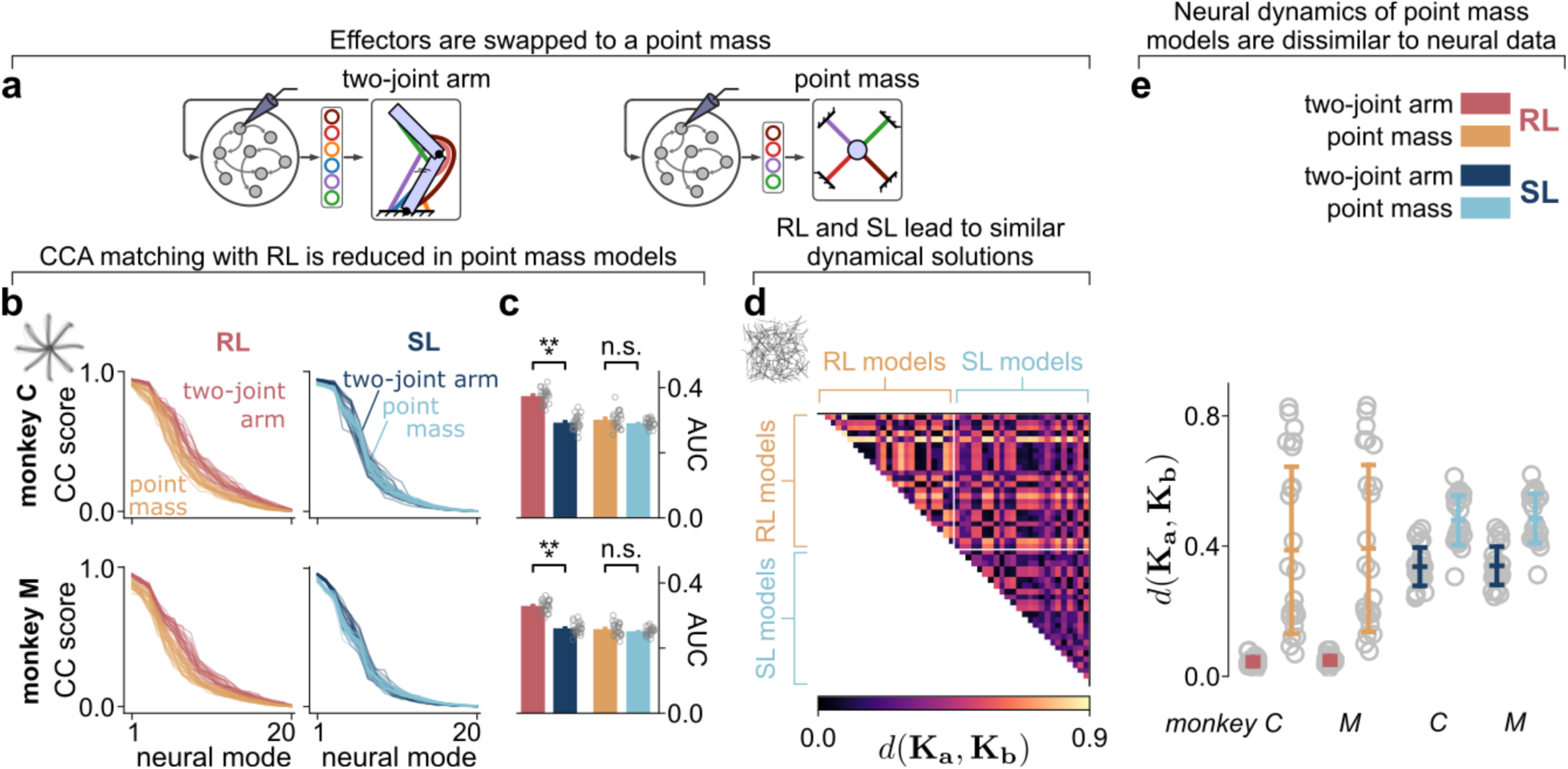
Simplifying biomechanical complexity of the effector drastically reduced the match between RL neural networks and neural recordings. **a**. We swapped the effector from a two-joints, six-actuators planar arm with Hill-type muscles to a simple point mass with four linear actuators. **b**, **c**. Learning to control a point mass instead of a two-joints arm greatly reduced Canonical Correlation scores between neural recordings and RL models during a center-out reaching task (n=24 per group). AUC, area under curve. **d**. Dissimilarity scores were also much closer between RL and SL models in a random target task. **e**. Dissimilarity scores against neural recordings indicate a collapse of similarity between neural recording and model dynamical solutions trained to control a point mass. Bar height and error bars indicate mean and standard deviations, respectively.

The effect was removed when comparing the arm model to a reduced arm model with 4 muscle actuators instead of 6, to match the number of actuators of the point mass (Fig. S10; rank-sum test, monkey C: U=-26.8, p<0.001 for both monkeys), confirming that the brain-like similarities result from the effector mechanics and not strictly the dimensionality of the control scheme.

We then applied DSA on neural activity of point-mass models during random-to-random reaches. Unlike networks driving the biomechanical arm model, RL- and SL-trained networks had similar Procrustes dissimilarity scores (Fig. 5d-e, U=-1.52, p=0.13 for model-model comparisons; T=88, p=0.08 for both monkeys in model-recording comparisons), indicating the control solutions are computationally similar. This convergence to dynamically similar solutions could be attributed to a flooring effect when controlling such a dynamically simple actuator. More broadly, this demonstrates that under certain conditions, particularly pertaining to the level of complexity of control dynamics, learning rules may yield universality of dynamical solutions^55^, but this universality property can break down under more realistic settings. Finally, we applied this analysis to neural recordings of monkeys performing a random target reaching task and observed that RL- and SL-trained models were much more dissimilar to neural recordings when trained to control a point-mass effector than when they were trained to control an arm. Broadly speaking, our results suggest that the biomechanics of the effector constrain the computational solution expressed the circuits. This emphasizes the importance of developmental trajectories on the computational solutions achieved by a network: RL-trained networks align well with primate motor cortex, but only if they are optimized for a primate-like arm.

### Reinforcement learning and supervised learning shape different dynamical regimes of neural activity

Since RL-trained models yield dynamics that are closer to those displayed by neural recordings, we sought to understand the nature of these dynamics. We leveraged our simulated networks to gain insight into neural computation which would otherwise be unavailable in experimental data. A compact and theoretically well-defined way to characterize the dynamics of a system is to identify fixed points in state space where activity does not change over time^44,45,56^ (*i.e.*, ***ḣ*** = **0**, see Methods). Fixed points have been shown to be important dynamics features for a number of tasks^44,56^, where their stability properties can reveal a great deal about learned solutions. When autonomous network dynamics contain stable fixed points, they are readily obtained by simulating the networks forward in time until its activity settles to that fixed point in neural space. By varying initial conditions (or applying perturbations), one can study subsequent trajectories toward the fixed points. This allows us to discriminate between three possible scenarios: (i) that perturbations onto the initial states are quickly resorbed over time, producing a single stable fixed point (contractive dynamics; Fig. 6a, upper illustration); (ii) that they are amplified (expansive/ chaotic dynamics; lower illustration); or (iii) that their mutual distance remains similar over time and settle into individual fixed points whose structure reflects the initial perturbations (neutral or isometric dynamics, middle illustration). Each of these dynamical regimes display distinct computational advantages. Contractive dynamics make dynamical systems robust against perturbations because neural trajectories easily recover to the same fixed point, but trajectories become stereotypical and struggle to retain information from past states. Expansive dynamics naturally maintain a memory trace of previous neural states because they do not collapse over time to a common state but are unstable against neural noise and perturbations. Isometric dynamics, often referred to as the “edge of chaos”, combine the above benefits while avoiding the associated caveats^56,57^.

**Fig. 6:**
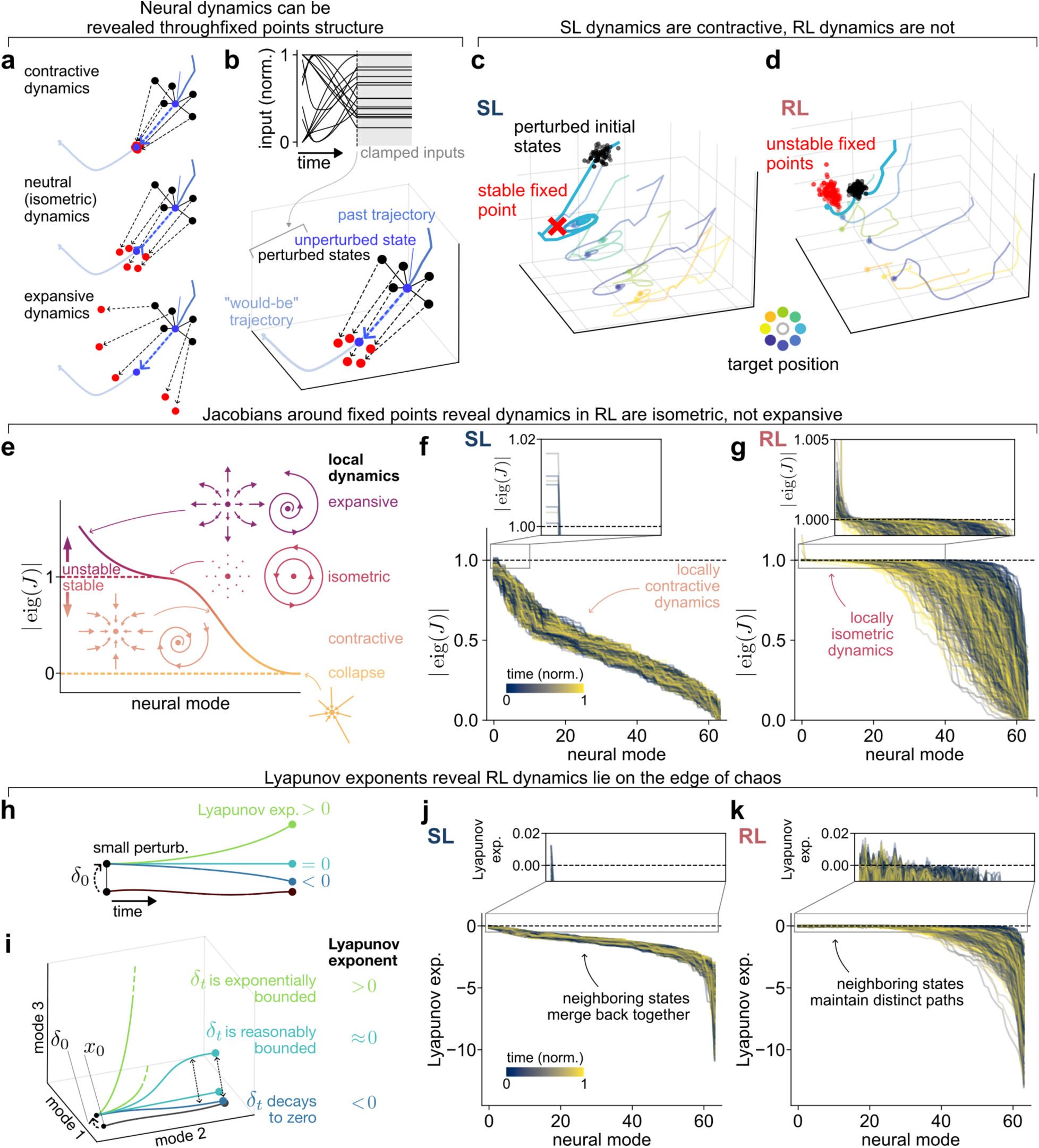
RL models produce dynamics that lie on the edge of chaos. **a.** Network models could be ruled by a range of dynamics, such as contractive or expansive dynamics, but also non-contractive, non-expansive dynamics (“isometric”). **b.** To test between these possibilities, fixed points were computed across all timesteps by perturbing neural states, fixing input values at that time and running forward passes until hidden activity settled to a fixed point (see Methods). **c-d.** Example neural trajectories, perturbed initial states and resulting fixed point(s) for a SL model (c) and RL model (d). **e.** The eigenspectrum of the recurrent activity’s Jacobian of a model can inform the (locally linear) dynamics around a given fixed point. **f-g.** Modulus of the Jacobian eigenspectrum at the unperturbed fixed points across all SL models (f) and RL models (g). Each curve is color-coded based on time within a single reach, with 0 in the color code being the start of the reach and 1 being the end of the reach. **h.** Lyapunov exponents can be computed by quantifying the rate of divergence between an original trajectory (black line) and a trajectory unfolding from an initially close-by state obtained with a small perturbation *δ*_0_. In different colors are represented trajectories that would yield a negative, null, or positive Lyapunov exponent. These indicate a trajectory that converges back, maintains a distance close to *δ*_0_, and diverges away, respectively. **i.** Non-negative Lyapunov exponent enable new trajectories, such as moving into new dimensions in state space, generally improving the expressivity of the dynamical system. If these new trajectories are reasonably (non-exponentially) bounded, their Lyapunov exponents will remain close to zero, providing expressivity without explosive instability. **j-k.** Lyapunov spectrum for each fixed point resulting from forward passes on the unperturbed initial state, for all SL models (j) and RL models (k).

Similar to motor cortices^58^, our model dynamics heavily depend on constantly changing inputs, making them non-autonomous and preventing us from computing fixed points as-is. Nevertheless, for moderately changing inputs, one expects the stability profiles of the systems to remain. To analyse dynamics regimes, we clamped the input vector at a given time *t* and ran the models autonomously until a fixed point convergence criterion was met (Fig. 6b, see Methods for details), thereby obtaining an estimate of “input-dependent” fixed point. We also performed this procedure after we applied small perturbations to the initial neural states from and repeated the forward passes to observe how neural dynamics varied from neighboring neural states. (Fig. 6b). This approach allowed us to assess the impact of learning rules on the stability of the dynamics.

Overall, SL-trained models displayed strong convergence of neural activity across initial states to overlapping fixed points, suggesting these optimized models tend to operate in a contractive dynamical regime (Fig. 6c). In contrast, RL-trained models tended to settle onto non-overlapping states, producing a coherent cloud of fixed points (Fig. 6d), meaning that RL-trained models could be expansive or isometric. To formally investigate this, we inspected the eigenvalues of the RNN’s forward map’s Jacobian evaluated at the fixed points (Fig. 6e). The eigenspectra of the Jacobian matrix for unperturbed fixed points were radically different in SL- and RL-trained models, the former being much shallower with barely any leading eigenvalues with modulus greater than one (Fig. 6f), while the latter produced many eigenvalue moduli close to—although generally below—one (Fig. 6g). This suggests that network dynamics were non-contractive and non-expansive in RL-trained models, indicative of an isometric, marginally stable dynamical system. This difference in dynamical regime arose despite identical initialization schemes and initialization seeds (see Methods, *Learning algorithms* section for details).

A more formal proposal is that RL-trained models lie properly at the edge of chaos^59^, a dynamical regime at the interface between order and chaotic regimes. This is more stringent than having the Jacobian of a discrete-time map with eigenvalues whose modulus close to one because beyond the local dynamics around the fixed points, it informs us about the general expressivity–and therefore, computational potency–of the neural dynamics it produces over longer timescales^47^. Intuitively, if two close-by neural states produce different, diverging trajectories, the number of distinguishable trajectories our network can produce increases accordingly. Mathematically, this is defined as a system with Lyapunov exponents (the exponential rates of divergence between infinitesimal perturbations; Fig. 6h) that are positive^60,61^. However, such a system is explosively unstable (chaotic) which can be computationally brittle. Rather, trajectories whose distance is non-exponentially growing will maintain this expressivity without suffering from instability (Fig. 6i). Such system will have Lyapunov exponents close to zero. Note that both types of systems can produce locally unstable dynamics around a fixed point if the latter stabilizes further away from the fixed point, which will not be apparent from analysis of the fixed point alone.

We computed the Lyapunov spectra of our RNN’s forward maps around the unperturbed fixed points found above and observed a stark difference between SL- and RL-trained models. Reminiscent to the eigenvalue of the Jacobians around the fixed points, the Lyapunov spectra stayed close to zero in the RL-trained models but not in the SL-trained models (Fig. 6j-k). This suggests that RL-trained models tend to evolve at the edge of chaos. By lying at this interface, such systems avoid the caveats of chaotic instability with several advantages. They express a richer repertoire of dynamical motifs^47^, retain a more sustained memory trace of previous activity over long timescales^47,57,62^, and as a consequence more easily maintain loss gradients that neither vanish nor explode, accommodate smoother plasticity during continual learning^48,57,63^. We note that a growing body of work argues that neural activity in brains lie at the edge of chaos (for a recent review, see Ref. ^59^).

### Reinforcement learning provides more flexible neural dynamics for subsequent adaptation to perturbations

We lastly tested whether the edge of chaos regime truly allowed RL-trained networks to more smoothly learn and generalize. While the *de novo* learning modeled above developmentally shapes neural dynamics, animals continue to acquire new skills throughout their life. It is advantageous during early developmental stages to yield neural representations that enable rapid and continuous learning through the lifetime of the animal. We next tested whether RL-trained networks demonstrate a brain-like capacity for continuous adaptation. A well-characterised example of this phenomenon is sensorimotor adaptation, where individuals learn to recalibrate the mapping between sensory inputs and motor outputs based on predictable changes in the environment^49,64^. Experimentally, this can be studied by introducing a visuomotor rotation between the hand position and the visual feedback when reaching to a target (Fig. 7a). Behavioral experiments in both monkeys and humans demonstrate that the brain can counteract this mismatch between motor output and sensory feedback over repeated reaching attempts^8,49,64–66^.

**Fig. 7:**
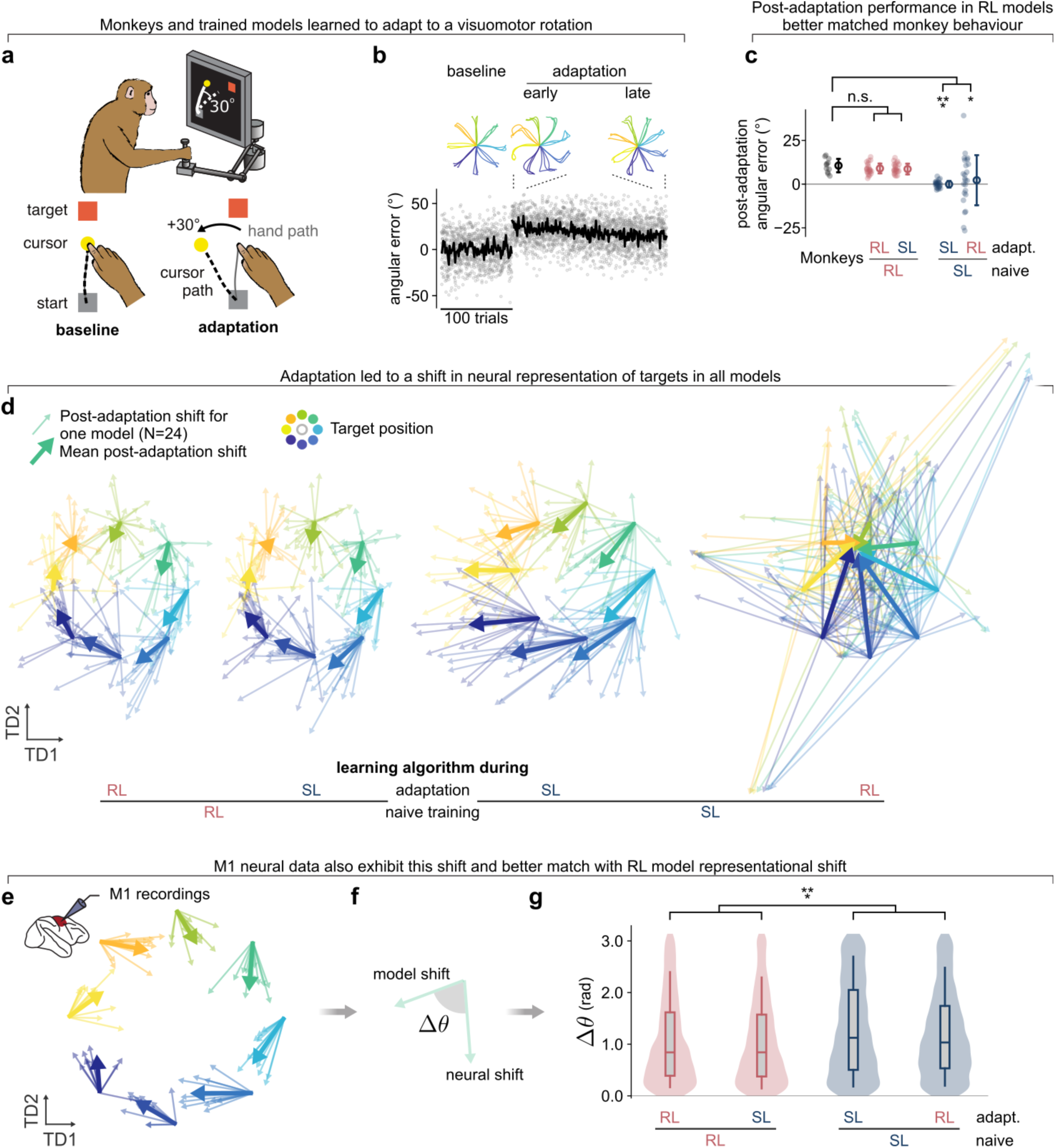
Exposure to visuomotor perturbations produce biologically plausible neural reorganization only in RL-trained models. **a**. Monkeys were exposed to 30° visuomotor rotations while performing a center-out reaching task. **b**. Mid-reach angular deviation from a straight trajectory across trials for both monkeys over 11 sessions. The thick black line indicates the mean across sessions. **c**. Angular deviation averaged over the last 8 trials of the adaptation block for each monkey session (black) and over thirty-two simulated trials post-adaptation for each neural network (n=24 seeds per group). The open circle and error bars indicate the mean and standard deviation, respectively. **d**. Shift of neural network activity in the target-predictive TDR subspace with arrows pointing from pre-to post-adaptation decoded states. Thick arrows represent across-session vector means. **e**. Same as panel d for neural recordings across both monkeys and 11 sessions. **f**, **g**. We quantified the similarity between neural networks and neural recordings as the angular difference Δ*θ* between neural shifts in TDR subspace. TDR: Targeted Dimensionality Reduction.

We exposed the monkeys to a 30° visuomotor rotation while recording from the motor cortex. We found that the animals counteracted the perturbation over tens of trials, gradually straightening their reach paths before achieving asymptotic performance with a small residual error (Fig. 7b). We quantified their performance by measuring the mid-reach angular deviation from a straight line of trajectories at the end of adaptation. (Fig. 7c). We then exposed the 24 seed-matched neural network models of each type (RL and SL) to the same behavioral perturbation. We tested the ability of each learning rule to adapt to the visuomotor rotation, explicitly considering the possibility that this continuous adaption need not employ the same learning rule as *de novo* development^65,66^. This gave four possible scenarios: *de novo* learning on RL and subsequent adaptation on both RL and SL, plus *de novo* learning on SL and subsequent adaption with both RL and SL.

Curiously, the asymptotic angular error of the monkeys was similar to that of models trained *de novo* with RL regardless of the learning scheme used for adapting to the perturbation (Fig. 7c; rank-sum test, RL-RL: U=1.10, p=0.27; RL-SL: U=0.82, p=0.41). In contrast, we saw striking differences in the behavioral performance of models trained *de novo* with SL (SL-SL: U=4.65, p<0.001; SL-RL: U=1.99, p=0.04). When SL was also used for adaptation (SL-SL), the models over-learned the visuomotor perturbation compared to monkey behaviour. Surprisingly, the highest error occurred when RL was used for subsequent adaptation after SL pre-training (SL-RL). These differences highlight that the choice of learning rule can dramatically influence the ability to perform subsequent adaptation, and that networks trained *de novo* from RL produce the most monkey-like output. Analysis of model performance at different adaptation epochs show that this result was not dependent on duration of adaptation training (Fig. S12).

To better understand the causes of this behavioural discrepancy, we analysed changes in the neural representations throughout sensorimotor adaptation in each network models. Previous studies of similar motor adaptation in primate motor cortex demonstrates that neural activity “shifts” in an orderly fashion^49,64^, with neural states encoding for a given target orientation essentially rotating along a circle formed by the pre-adaptation target encoding states (Fig. 7e).

Given the behavioral (Fig. 7c) and dynamical (Fig. 4c) similarities between RL-trained networks and monkeys, we hypothesized that only networks trained *de novo* with RL would show a similar reorganization. To test this hypothesis, we applied targeted dimensionality reduction (TDR) to each model’s pre-adaptation neural states and assessed the amount and direction of shift in these target-specific states after adaption. We found that all models trained *de novo* through RL (RL-RL and RL-SL) showed such an orderly shift in neural states (Fig. 7d), in agreement with the motor cortical data. In contrast, models trained *de novo* with SL showed a variety of behaviour depending on the learning rule used for subsequent adaptation. If SL was applied during adaptation (SL-SL), we observed a full translation of neural states as opposed to a rotational shift, suggesting that the networks learned a whole new representational structure. If RL was applied during adaptation (SL-RL), the representational structure completely reshuffled, resulting in no consistent shift across model seeds. We interpret this high-variance restructuration as an indication that the neural representations learned during SL are challenging to leverage for RL-driven learning schemes, forcing a complete re-acquisition of a control policy rather than simply adapting the original control policy to the new, perturbed dynamics of the environment.

To quantify these differences across models, we compared the angular difference between shifts in neural states of our models to the shifts observed in M1 across 11 sessions for 2 monkeys during visuomotor adaptation (Fig. 7e-f). Indeed, this distribution of angular differences was significantly smaller for RL models compared to SL models but was equivalent for RL models which adapted to the perturbation using RL or SL (Fig. 7g; RL-RL vs SL-SL: U=-7.94; RL-RL vs SL-RL: U=-5.75; RL-SL vs SL-SL: U=-8.40; RL-SL vs SL-RL: U=-6.09; p<0.001 for all comparisons). Together, these results show that networks that acquire their *de novo* behavioral repertoire through RL show brain-like properties during subsequent adaptation and learning, both behaviorally and neurally.

## Discussion

By comparing *in silico* neural representations from two putative learning rules to neural recordings of monkey motor cortex, we demonstrate that the neural control solution expressed in biological systems is more aligned with a learning process dominated by RL from both a geometric and dynamical perspective, as opposed to a direct error-minimizing SL scheme. We also demonstrate how the biomechanical complexity of the control system enables this discriminability in neural solutions under each learning scheme. Dynamical analyses revealed that our RL-trained networks produce activity that, unlike their SL-trained counterparts, operate at the edge of chaos, which is well-known to provide computational and plasticity advantages. They maintain a trace of previous neural activity without contracting back to fixed points, which enables better memory capacity. This also allows for better transport of credit assignment over time, preventing vanishing gradients that could lead to representational collapse during continual learning^63^. Consistent with this possibility, we showed that RL, unlike SL, yields neural representations that allow for smooth shifts in encoding states through continual learning during sensorimotor adaptation. These analyses allowed us to distinguish between two well-established hypotheses on the nature of a dominant teaching signal for neural learning.

There is a rich history of comparing RL and SL algorithms and their relevance to neural learning^67–69^. Rueckauer and van Gerven (2020)^67^ show that RNN models can be trained equivalently using SL and RL models. Song *et al.* (2017)^69^ elegantly demonstrate that RL-trained RNNs can learn solutions from value-based decision-making tasks that qualitatively match empirical observations. Our study expands these findings to neural responses in motor cortex, and more importantly, directly confronts the modeling results to matching neural data, bridging the gap between modeling and experimental works. More recently, Aldarondo *et al*. (2024)^68^ used a clever autoregressive approach and imitation learning (a special type of RL) to train models to reproduce behavior from freely behaving rats. Consistent with our findings, this RL approach outperformed a SL approach in predicting recorded neural activity in the dorsolateral striatum and motor cortex. While this powerful method (learning through imitation) relies on an existing corpus of pose estimation data, we demonstrate that these results hold in a more ethological generative setting (learning through action).

The current study hinges on using the same dense reward and loss formulation across RL and SL models to enable meaningful comparisons between learning schemes. This departs from prevailing views in systems neuroscience that RL’s advantage lies in its ability to tackle sparse and delayed rewards^21,32,69^. Pioneering work bridging RL and systems neuroscience heavily focused on cognitive and decision-making tasks^21,22,26,32–34^, for which reward patterns usually match these characteristics. We believe this enabled early progress, as delayed rewards could easily be dissociated from other confounding factors experimentally. In contrast, the sensorimotor domain usually deals with dense signals, and indeed most recent work on this topic employs dense, not sparse, reward functions^16,24,43,70–75^. We argue that RL’s usefulness lies in its lack of initial assumptions about the relationship between action and outcomes, instead relying on directly “learning from experience” (or as it was originally put by Sutton and Barto, “learning from interaction”^17^). Indeed, this is what allows it to gracefully deal with sparsity or the lack of temporal contingency, while also overcoming the long-standing issue of non-differentiability in the motor system.

Learning in the central nervous system encompasses many partially overlapping mechanisms occurring at different spatial, temporal, and neurobiological scales^5,7,8,19,49,76,77^. For instance, decision-making involves learning the correct set of choices to make when faced with a series of behavioral options^21,32,33^, while sensorimotor adaptation entails remapping sensory and motor primitives to match small-scale but consistent changes in environmental dynamics^65,66^. Here we focus on *de novo* learning, which is learning from scratch, such as when a newborn first interacts with the world^18–20^. In the context of neural control of movement, we draw an analogy to the acquisition of a repertoire of basic motor actions, such as moving from one arbitrary point in space to another^78^. Such actions can then be combined into complete movements through sequence learning^76,79^, and then into complex, purposeful motor skills through cognitive modulation of whole-body actions. However, these later learning stages rely on the existence of the rich, well-tuned repertoire of core movement representations built early in an organism’s lifespan^18^.

It is important to emphasize that the dominant teaching signal likely varies across these intertwined processes. For instance, sensorimotor adaptation is driven by a process akin to supervised learning^8,9,65^. In this light, our adaptation experiment showcases a central point on the interplay between learning processes: that neural representations resulting from one learning scheme can still provide fertile ground for another learning scheme to take over and produce purposeful plasticity, building on previous representation through ordered reorganisation rather than remodelling it entirely. Even during *de novo* motor learning, which this study focuses on, unsupervised learning captures a significant portion of developmental plasticity in biological systems^80,81^ and demonstrably relates to the somatotopic organization of sensorimotor cortices across species^82^. It is also a viable means of enforcing efficient state space exploration in over-actuated systems like natural musculoskeletal effectors^83^. While our models do not include spatial connectivity arrangement, we note that this is an advantage in pursuing our main finding as our models’ fully connected architecture alleviates topographic constraints. Nevertheless, the co-occurrence of neural plasticity driven by unsupervised learning in the brain illustrates our point that several learning schemes likely co-exist but unevenly shape and constrain neural activity^5,8^.

At a finer scale, a prominent topic under investigation is finding which learning rules govern plasticity mechanisms at the synaptic level in the brain^42,84,85^. Particularly, gradient descent is one such mechanism that is widely popular in machine learning as it is fast and efficient^86^. While its biological plausibility has been challenged, there is now mounting evidence that it could reasonably be approximated locally at the synaptic level, making it an appealing hypothesis to explain neural learning^10,87–89^. It is important to appreciate that both reinforcement and supervised learning, in the forms we use them here, rely on gradient descent, the former through the policy gradient theorem^16^ and the latter directly through explicit derivation of the loss^86^. Therefore, this work does not provide definitive evidence for or against the existence of gradient-based learning in the brain, but rather offers some insights on what types of gradients might be pursued by plasticity rules, namely those stemming from RL or SL schemes.

We view neural learning required for skill acquisition as a three-steps process (though note that other types of learning, such as cognitive decision making^90^ or sequence learning^76^, need not follow all three steps). First, a teaching signal provides a global performance metric^72,91^. Second, a credit assignment mechanism dictates the contribution of each synaptic connection in the circuit to this metric value^10,11,16,88^. Third, the strength of the synaptic connection is modulated according to an update rule and proportionally to that synapse’s estimated contribution to performance^89^. In this study, we propose and demonstrate that *de novo* learning at the neural circuit scale is likely dominated by a reinforcement-based teaching signal. In other words, once a value function is estimated, (potentially gradient-based) optimization is performed over that value function rather than a loss function directly. What credit assignment and synaptic update rules drive plasticity at the local circuitry level to give rise to such a coordinated process remain open and ongoing questions^10,89,92^. Particularly, how a reinforcement-based teaching signal would interact in practice with different plasticity and update rule hypotheses are exciting avenues of research for future studies.

## Acknowledgements

We thank Carol Massumoto for the monkey illustration. We thank Dr. Raeed Chowdhury, Dr. Jonathan A. Michaels, and Dr. Colin Bredenberg for their helpful feedback on early drafts of the manuscript. This work was supported in part by the FRQNT Strategic Clusters Program (Centre UNIQUE - Centre de recherche Neuro-IA du Québec, to O.C.), the Fonds de Recherche du Québec secteur Santé (FRQ) to the Centre Interdisciplinaire de Recherche sur le Cerveau et l’Apprentissage (to O.C.), NSERC Discovery Grant (RGPIN-2018-04821, to G.L.), Canada CIFAR AI Chair Program, Canada Research Chair in Neural Computations and Interfacing (CIHR, tier 2, to G.L.), and grant chercheurs-boursiers en intelligence artificielle from the Fonds de recherche du Quebec Santé (to M.G.P.).

## Author Contributions

O.C., G.L, and M.G.P. devised the project. O.C. and N.H.K. performed modeling and simulations. O.C. and N.H.K. analyzed data and generated figures. M.G.P. collected the monkey datasets. O.C., G.L., and M.G.P. wrote and edited the manuscript. G.L. and M.G.P. jointly supervised the work.

## Data & Code Availability Statement

All analyses and simulations were implemented with custom Python code and open-source software libraries (see Methods for details). Code to replicate all simulations and analyses will be available prior to publication, or sooner by reasonable request. The majority of sessions analyzed of the monkey datasets are already publicly available (https://dandiarchive.org/dandiset/000688).

## Methods

### Monkey behaviour

We trained two males *Macaca mulatta* monkeys aged 6–10 years to sit in a primate chair and make reaching movements using a planar manipulandum. The motion of the manipulandum handle was mapped to a cursor on a computer screen situated before the monkey (Fig. 7a) with the behavioural task being run through custom software in Matlab (The Mathworks). Monkeys were trained for at least several months to perform a two-dimensional center-out reaching task and a more complex random target sequential reaching task before the neural recording sessions to ensure they attained expert performance. For both tasks, we recorded the position of the endpoint at a sampling frequency of 1 kHz using encoders in the joints of the manipulandum. Portions of the center-out reaching data have been previously published and analysed in Refs. ^3,49,93–95^. Portions of the random target data have been previously published and analysed in Refs. ^3,96,97^. We briefly summarize the task paradigms below.

In the center-out task, the monkey moved its hand to the center of the workspace to begin each trial. After a variable waiting period, one of eight targets was displayed. All targets were equally spaced in a circle and selected randomly with uniform probability. A go cue sound signalled the monkeys to initiate a reach to the target. They were required to reach the target within 1 s after the go cue and hold inside the target for 0.5 s to receive a liquid reward. We included eighteen and six experimental sessions from monkey C and M, respectively, for the center-out reaching task. The duration of each session varied from 154 to 217 successful reaches as monkeys reached for as long as possible to ensure behaviour had sufficient time to stabilize^49^. For eight and three sessions for monkey C and M, respectively, the monkey was then exposed to a follow-up adaptation block with a counter-clockwise 30° visuomotor rotation between the onscreen cursor position and veridical position of the manipulandum. Monkeys performed 159-217 successful reaches during baseline and 243-336 successful reaches during adaptation blocks. The last 100 successful baseline trials and first 243 successful adaptation trials for all sessions are displayed in Fig. 7b.

In the random target task, the monkey made four consecutive reaches to random targets within a 10×10 cm^2^ workspace in each trial. Each target was presented sequentially in a random location within a 5 cm to 15 cm distance from the previous target to ensure a consistent reaching distance range. There was no go cue and the monkey had to hold the cursor inside the target for 100 ms before the next target was presented following a 100 ms delay period. The monkey received a liquid reward during a short break after each successful sequence of four target acquisitions. For this study we restricted analysis to the first reach from each four-reaches sequence to avoid any effect of sequence order.

### Neural recordings

All surgical and experimental procedures were approved by the Institutional Animal Care and Use Committee of Northwestern University under protocol no. IS00000367. We implanted 96-channel Utah electrode arrays in the primary motor cortex (M1) and dorsal premotor cortex (PMd) using standard surgical procedures. Implants were positioned in the hemisphere opposite to the hand animals used to perform the task. Monkeys C and M received an array in M1 and PMd simultaneously. Neural activity was recorded with a Cerebus system (Blackrock Microsystems). The recordings on each channel were band-pass filtered (250 Hz–5 kHz) and then converted to spike times via threshold crossing set to 5.5 times the root-mean-square activity on each channel. Spikes were manually sorted to identify putative single neurons. For the center-out reaching sessions, the average number of units included was: monkey C, 116.5 ± 91.3 (mean ± s.t.d.; range, 55–353); monkey M, 89.0 ± 31.8 (range, 52–129). For VR sessions, we only included M1 recordings, leading to: monkey C, 70.3 ± 10.6 (range, 60–95); monkey M, 44.3 ± 5.0 (range, 39–51). For the random target sessions, the number of units recorded were 295 and 163 for monkey C and M, respectively. A more detailed description of the behavioural and neural recording methods is presented in Ref. ^49,93^.

### Models

#### Architecture

All neural network models were implemented using the open-source Python package pytorch^98^. Models consisted of a recurrent neural network (RNN) composed of gated recurrent units (GRUs) with *tanh* non-linearity according to the equation below.

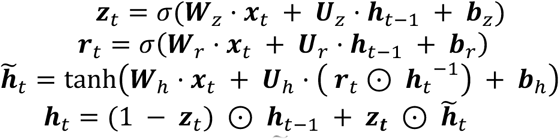

The symbol ⊙ represents the element-wise product, ***h̃***_*t*_ the candidate hidden state at timestep *t*, ***h**_t_* the hidden state at timestep *t*, *σ* the sigmoid function, *U_z_*, *U_r_*, *U_h_*, ***W**_z_*, ***W**_r_*, ***W**_h_* are learnable weight parameters, and ***b**_z_*, ***b**_r_*, ***b**_h_* are learnable bias parameters. The GRU layer then projects into a fully connected output layer of six neurons (Fig. 2a-b) and a sigmoid non-linearity to ensure positive outputs. Each output from this last layer served as an action to be passed to one of the six actuators controlling the effector. In the case of the point mass models, the output layer contained four neurons as the point mass was controlled by four actuators (Fig. 5a). The RNNs were composed of sixty-four GRUs, although our results remained similar across a range of sizes (Fig. S3). They received as input a vector containing a step go cue signal, the cartesian position of the effector’s endpoint, the start position and target position, as well as each actuator’s length and velocity^43^ (Fig. 2c). For the RL models, this input vector was also provided to a separate critic network, which consisted of a multi-layer perceptron containing four layers of 100, 64, 64 and 1 neurons, with the second and third layers including a *tanh* non-linearity. The output of the perceptron served as the value estimate for training the RL models.

#### Environments

The environment models employed were from the open-source Python package motornet^43^. Specifically, we employed the *RigidTendonArm26* effector with default parameter values and the *Runge-Kutta 4* algorithm for numerical integration. The actuators were *MujocoHillMuscle* object with no passive force contribution and *l_min_* = 0.3, *l_min_* = 1.8. For the point mass simulations, we employed the *ReluPointMass24* effector and set its mass to be equal to the combined mass of both arm segments (1.82 + 1.43 = 3.25 *kg*). The point mass was actuated by a *ReluMuscle* object with time constants set to the same values as the *MujocoHillMuscle* (*τ_act_* = 0.01, *τ_deact_* = 0.04) and maximal isometric force of 1000 N. The point mass actuators were positioned in a “X” configuration, with each anchor point coordinates being [±4, ±4] m and the origin being the center of the 2×2 m space in which the point mass was evolving. The simulation time constant was set to 10 ms for all models.

#### Training procedure

All models were trained to reach from a random start position to a random target of 1 cm radius drawn uniformly within the full joint space. Actuator endpoints were required to stay inside the start position until the go cue signal at 200 ms, and to go to the target afterwards. Episodes lasted 5 s and were terminated early if the effector endpoint remained inside the target for 800 ms consecutively. The instantaneous reward for RL models was:

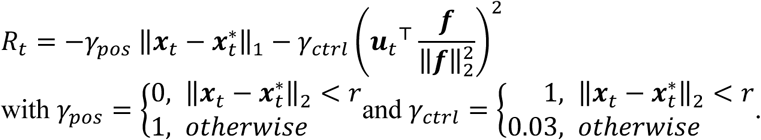

***x**_t_*, 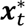, ***u**_t_*,***f*** were the position, target, action, and maximum isometric contraction vectors used for normalization of the control penalty, respectively. Therefore, the theoretical upper bound of *R_t_* was 0. The instantaneous loss for SL models was simply the negative of the reward function.

During adaptation to visuomotor rotations, the input weights were updated over training epochs while all other weights remained frozen to match previous work^49^. To ensure that freezing all weights except input weights did not single-handedly explain our results, we repeated the same adaptation experiments with plastic recurrent weights and show that our findings remain consistent (Fig. S13).

#### Learning algorithms

Both RL and SL used a learning rate of 3e-4, gradient norm clipping of 0.5, and a batch size of 128. We collected 128 simulation steps per rollout before performing a backward pass and network weight update. In both cases we applied independent Gaussian action noise *ε*∼𝒩(0, *σ*^2^) with periodical re-sampling every *n*∼𝒰{*a*, *b*} timesteps (*a* = 16, *b* = 24) to smooth out exploration^73^. We also added a L2 regularization on neural activity and its derivative to the gradient for both model groups to promote parsimonious and sparse network activation levels, which usually leads to more biologically plausible activity patterns^99^. Regularization weights were *γ* = 0.01 and *γ* = 0.1 for neural activity and its derivative, respectively. Note that these regularization values did not impact the differences in Canonical Correlations between model groups (Fig. S1 and S2). Models were initialized with orthogonal initialization with a gain of 2 for input, recurrent, and output weight matrices. Seeds (n=24 per group) were matched between groups to ensure any difference between models did not arise from differences in initial learnable parameter values. Biases were all initialized at 0 except for the output layer, whose bias was initialized at −10. This negative bias ensures that on initial training epochs, output activity is falling well into the low saturating part of the sigmoid non-linearity, leading to action vectors ***u**_t_* with a lower norm. In practice this greatly helps learning by promoting a stable initial regime of activity^43,75^.

RL models were trained using Proximal Policy Optimization^50^ with the critic being a separate network from the policy (Fig. 2b) to prevent learned value representations from mixing up with the computations learned for control of the effector and ensure meaningful comparison with SL models. We used an implementation of PPO derived from the stable baselines 3 package^74^ (version 2.0.0a13) modified to enable recurrent architectures for the policy network. All hyperparameters were left at default except for the batch size of 128 to match SL models’ batch size. The critic network weight matrices initialization was orthogonal with a gain of 1, and bias was initialized at 0 for all layers.

We used exploration annealing for RL models to promote action space exploration in early training and exploitation in fine-tuning stages of training. Annealing was done based on the mean return *r̂* of the past 20 rollout according to a sigmoid mapping so that the standard deviation of the Gaussian noise term was *σ*^2^ = 0.1/(1 + *e*^5+15·r̂^). Therefore, we have 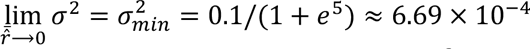. Although not required for learning, action noise was added to SL models and set to *ε*∼𝒩(0, 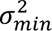) to match levels of action noise experienced by RL networks through most of training. While RL networks were exposed to higher action noise at the start of training, the mean return history *r̂* quickly dropped to low values (Fig. 2c) and most of training was spent in fine-tuning, ensuring RL models were experiencing action noise close to *ε*∼𝒩(0, 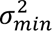) as well.

### Data analysis

All the analyses were implemented in Python using open-source packages such as numpy, matplotlib, scikit-learn and scipy^100–103^ and custom code. As we were analysing pre-existing datasets, no explicit planning of sample size, group randomization or blinding was performed. In all the analyses, we only included trials in which the animal successfully completed the task within the specified time constraints and received a reward.

#### Kinematics

We defined movement start and target acquisition times in a similar way in animal and model kinematics when compared to each other. Movement onset was defined as the last time radial velocity crossed a threshold of 3 cm/s before exiting the start position. The target was considered acquired 250 ms after the cursor’s distance to the target center was less than 1.5 cm and radial velocity was less than 4 cm/s. Center-out reach kinematics were interpolated to 100 timepoints between movement onset and target acquisition. For both model and monkey kinematics, angular error for trials with a visuomotor rotation was obtained at 150 ms after movement onset by computing the angular deviation of cartesian position from a straight line between the start and target position.

#### Neural data processing

Spike events from neural unit’s recordings were binned with a 10 ms bin size, square root transformed to stabilize variance^104^, and then smoothed using a non-causal Gaussian kernel with width *σ*^2^ = 50 ms to produce firing rate signals. Centre-out reach kinematics were interpolated to 100 timepoints between movement onset and target acquisition. For the visuomotor adaptation task, we employed a non-causal Gaussian kernel with width *σ*^2^ = 100 ms.

#### Statistical analyses

All statistical tests performed were two-sided. When indicated graphically, statistical significance was reported throughout with one, two, and three asterisks signifying a p-value of less than 0.05, 0.01, and 0.001, respectively. CC scores were computed on the leading 20 PCs of each model’s neural activity or pooled neural recordings from PMd (Monkey C: 192 recorded units; monkey M: 66 units) and M1 (Monkey C: 51 units; monkey M: 66 units), which explained 63% and 82% of neural variance for monkeys C and M, respectively. Using the first 10, 30, or 40 PCs in each dataset did not alter our main findings (Fig. S6). Principal component activity was column centered before computing canonical correlation. For comparison with biological recordings, we simulated as many trials in each target direction as was available in the neural recording session the simulations were compared to.

To compute dynamical dissimilarity scores, we simulated reaches with an identical start and end position as the monkey datasets during random target reaches. Like for CCA, DSA was performed on the leading 20 PCs of each model’s neural activity or pooled neural recordings from PMd (Monkey C: 211 recorded units; monkey M: 93 units) and M1 (Monkey C: 84 units; monkey M: 70 units), which explained 55% and 77% of neural variance for monkeys C and M, respectively. Dissimilarity scores were then computed on our dataset using the original study’s Python package^53^. Briefly, the Koopman operator was computed via dynamic mode decomposition^105^ on time-delayed embeddings of the neural data, which provides numerical stability while remaining diffeomorphic to the original system^106^. This approach corresponds to the Hankel Alternative View of Koopman (HAVOK) which is employed in dynamical systems analysis^107^. Finally, we computed the dynamical dissimilarity metric on the Koopman operators as a Procrustes dissimilarity over vector fields (Fig. S4a), which is a conjugacy-invariant version of Procrustes dissimilarity^53^. For this study, we retained the first twenty ranks for dynamic mode decomposition and constructed the time-delayed embeddings with ten delays. We selected these parameters by performing a hyperparameter sweep on neural data. Specifically, we selected the lowest number of ranks (to prevent over-fitting) to minimize reconstruction error on held-out data using a 10-fold cross validation procedure. Because we used bins of 10 ms for the neural data, 10 delays allowed us to capture dynamics spanning temporal ranges of at most 100 ms without drastically affecting reconstruction error (Fig. S14).

We performed targeted dimensionality reduction on neural data by fitting a Gaussian process regression model to predict planar target coordinates from neural data. The regression model was trained using the scikit-learn^102^ software implementation of algorithm 2.1 from Ref. ^108^ with normalization of target values and a composite kernel with radial basis function, constant and white noise kernel so that Cov(*x*, *y*) = *c*_1_ · 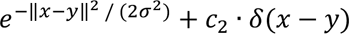. The optimization bounds for kernel parameters *c*_1_, *c*_2_, *σ* were left to default values. The model was fitted for each neural recording session on condition-averaged neural activity in the first 100 ms of movement in the last thirty-two trials of the baseline block and tested on the last thirty-two trials of the adaptation block. For simulation data, the regression model was trained on thirty-two reaches before adaptation and tested on the same number of trials after 500 epochs of adaptation. Our main findings remained similar when using 300 adaptation epochs, indicating that these results are at convergence (Fig. S12).

#### Fixed-point analysis

Fixed points are positions in neural state space for which the change in hidden activity ***ḣ*** is approximately zero (***ḣ*** = **0**) ^44,56^, or for a discrete system, when hidden activity at time *t* + 1 is equal to hidden activity at time *t* so that ***h**_t_*_+1_≈ ***h**_t_*. The existence and location of these fixed points depends on inputs ***u*** because ***h**_t_*_+1_ = *F*(***h**_t_*, ***u**_t_*), with *F* the nonlinear activation function of the network. Therefore, previous studies attempting to identify fixed points rely on static inputs^56^, while our models (and cortical areas such as M1 during movements^58^) employ dynamic inputs. To recreate similar conditions, we identified fixed points for every time points of our model simulations, clamping the input ***u**_t_* at time *t* over the optimization process. We performed forward passes on the trained recurrent networks with these constant inputs, starting with the hidden state ***h**_t_* or perturbed initial states from ***h**_t_* by adding Gaussian noise (*σ* = 0.05). For each time point *t*, we computed fixed points for the original unperturbed hidden state and 64 perturbed fixed points. This sampling around the original hidden state allowed us to determine the difference in flow field around the natural trajectory that models follow during evaluation. A fixed point was determined to exist when, for a given hidden state, when *q* < 10^-6^, for *q* = ½ (***h*** − *F*(***h***, ***u***))(***h*** − *F*(***h***, ***u***)). The optimization learning rate was reduced to 1e^-3^ to improve stability, and the maximum number of iterations has been extended to 80,000 to accommodate for the slower convergence. For the fixed point search, we employed the PyTorch implementation of the Fixed Point Finder package^109^.

Lyapunov exponents were computed using the standard QR decomposition algorithm^60^, starting from the hidden state at the unperturbed fixed point. Perturbations with Gaussian noise (*σ* = 0.05) were applied to this initial state to obtain 10 samples^61^. Each sample was then passed through the model for a hundred simulation timesteps. QR decomposition was applied to the Jacobian of the hidden state vector at each timestep to keep the directions of expansion orthogonal and avoid representational collapse into a few dominant modes, which is known to introduce numerical instability^60^. *Q* was initialized as the identity matrix. The diagonal elements of the upper triangular *R* matrix indicated the rate of expansion/contraction at that timestep. The Lyapunov exponent *λ*_*_for a neural mode *i* were computed as the temporal average of the log of each diagonal element:

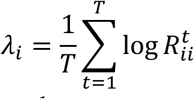

The resulting Lyapunov exponents are reported as a spectrum over neural modes *i*. Reported Lyapunov exponents at each timestep were the average value over the batch of perturbed initial states^61^.

---

**Fig. S1:**
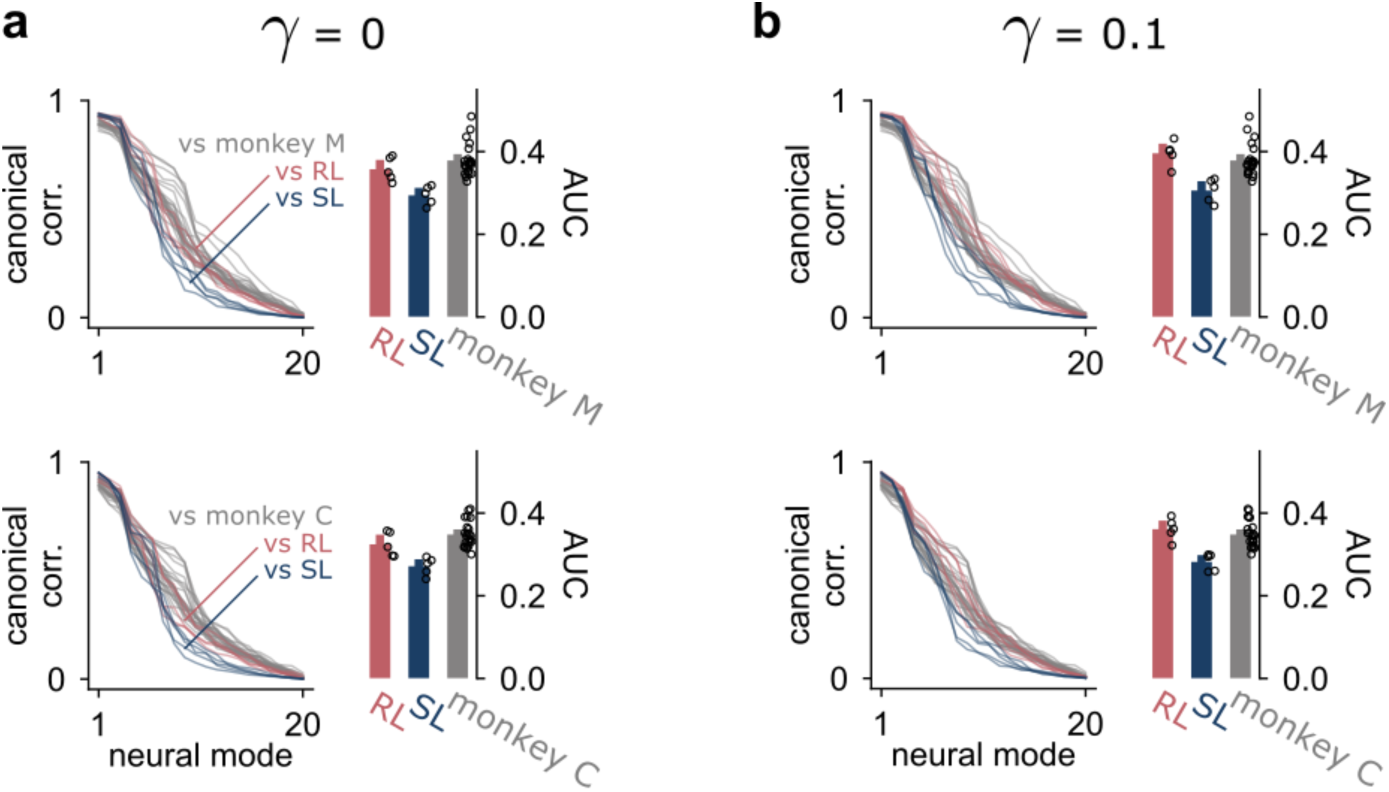
Varying the regularization weight on neural network hidden activation did not alter Canonical Correlation differences between learning rules. The default value was a penalty of 0.01. a. Regularization weight of 0. b. Regularization weight of 0.1. For each panel, the left subpanel shows the CC curves of five seed-matched networks against the same recording session for monkey C (top) and monkey M (bottom). The grey lines represent comparison to different recording sessions from the other monkey. The right subpanels show the area under curve (AUC) of the CC curves. Bar heights and error bars indicate the mean and 95% confidence intervals, respectively.

**Fig. S2:**
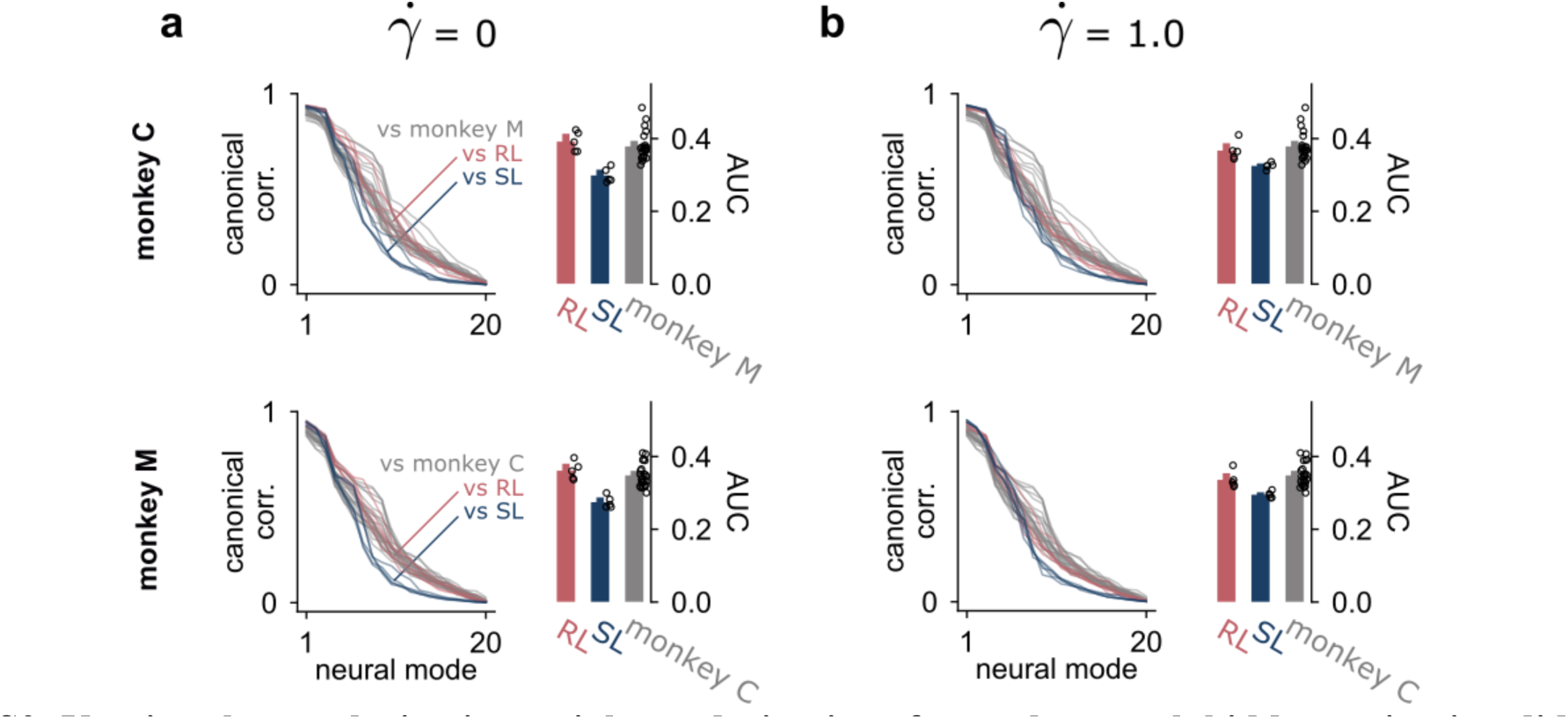
Varying the regularization weight on derivative of neural network hidden activation did not alter Canonical Correlation differences between learning rules. The default value was a penalty of 0.1 **a**. Regularization weight of 0. **b**. Regularization weight of 1.0. For each panel, the left subpanel shows the CC curves of five seed-matched networks against the same recording session for monkey C (top) and monkey M (bottom). The grey lines represent comparison to different recording sessions from the other monkey. The right subpanels show the area under curve (AUC) of the CC curves. Bar heights and error bars indicate the mean and 95% confidence intervals, respectively.

**Fig. S3:**
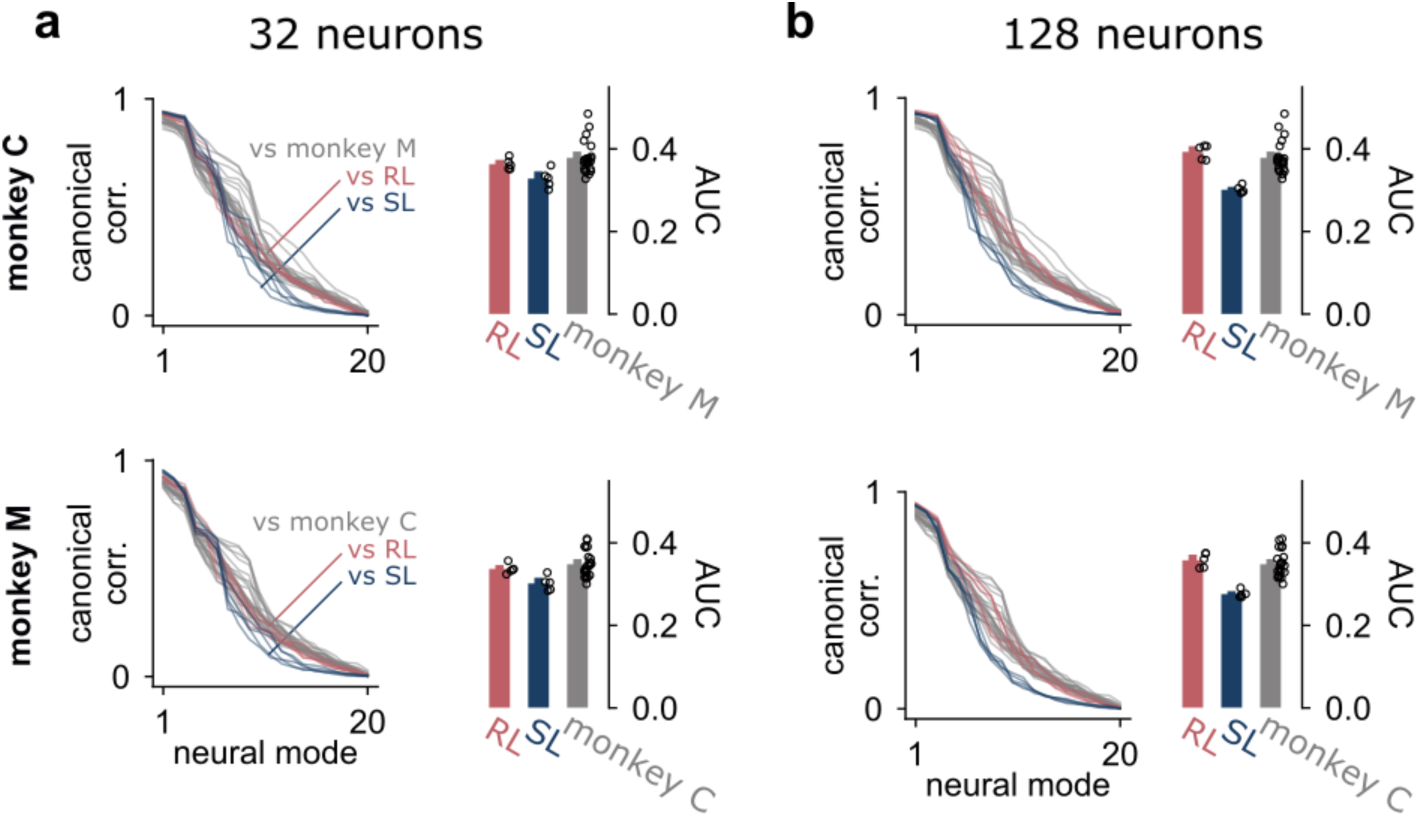
Varying the size of the neural network recurrent layer did not alter Canonical Correlation differences between learning rules. The default value was 64 neurons. **a**. Neural network size of 32 neurons. **b**. Neural network size of 128 neurons. For each panel, the left subpanel shows the CC curves of five seed-matched networks against the same recording session for monkey C (top) and monkey M (bottom). The grey lines represent comparison to different recording sessions from the other monkey. The right subpanels show the area under curve (AUC) of the CC curves. Bar heights and error bars indicate the mean and 95% confidence intervals, respectively.

**Fig. S4:**
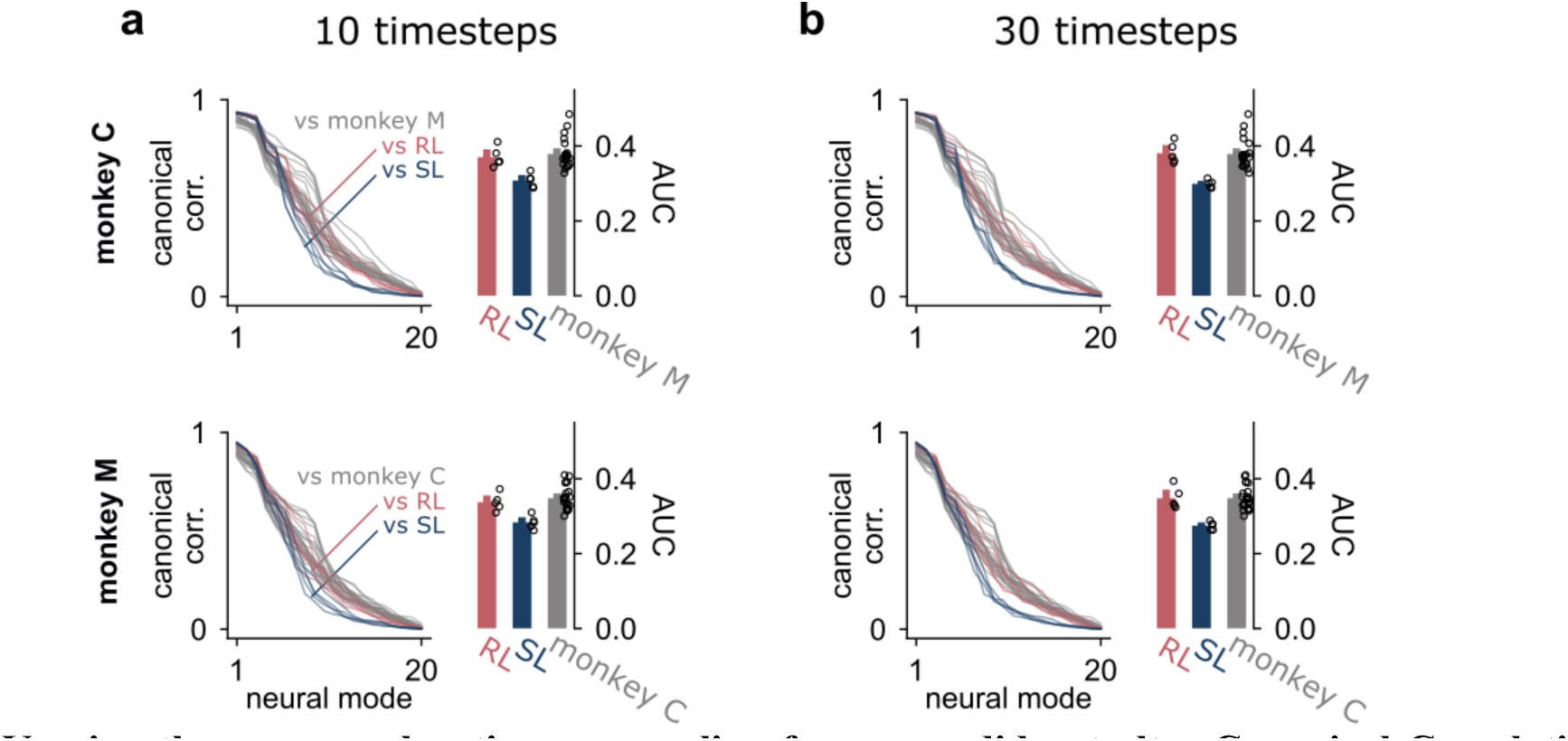
Varying the mean exploration re-sampling frequency did not alter Canonical Correlation differences between learning rules. The default mean frequency was 20 timesteps. **a**. Mean frequency of 10 timesteps. **b**. Mean frequency of 30 timesteps. **c**. Mean frequency of 40 timesteps. For each panel, the left subpanel shows the CC curves of five seed-matched networks against the same recording session for monkey C (top) and monkey M (bottom). The grey lines represent comparison to different recording sessions from the other monkey. The right subpanels show the area under curve (AUC) of the CC curves. Bar heights and error bars indicate the mean and 95% confidence intervals, respectively.

**Fig. S5:**
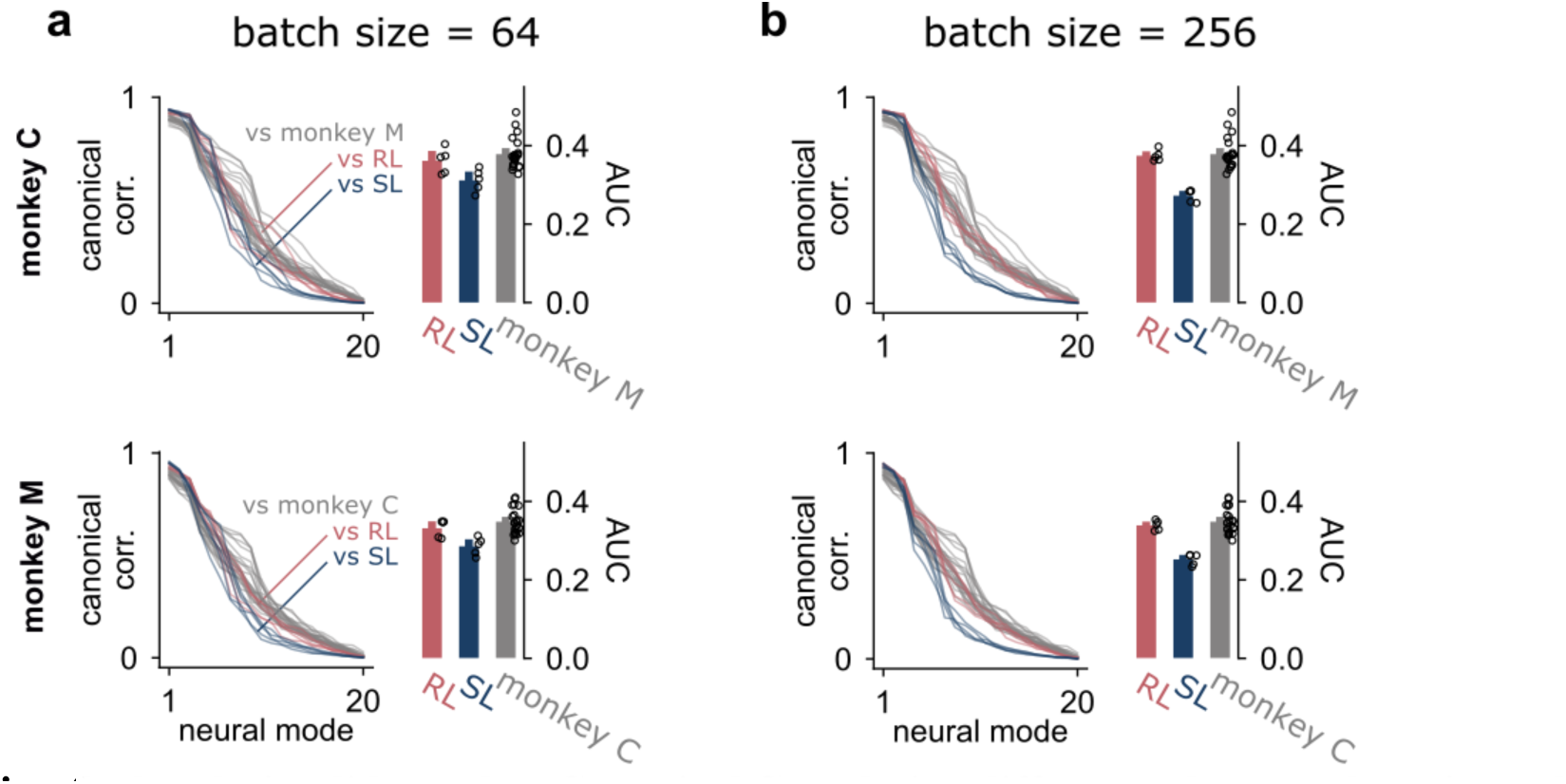
Varying the batch size did not alter Canonical Correlation differences between learning rules. The default batch size was 128. **a**. Batch size of 32. **b**. Batch size of 64. **c**. Batch size of 256. For each panel, the left subpanel shows the CC curves of five seed-matched networks against the same recording session for monkey C (top) and monkey M (bottom). The grey lines represent comparison to different recording sessions from the other monkey. The right subpanels show the area under curve (AUC) of the CC curves. Bar heights and error bars indicate the mean and 95% confidence intervals, respectively.

**Fig. S6:**
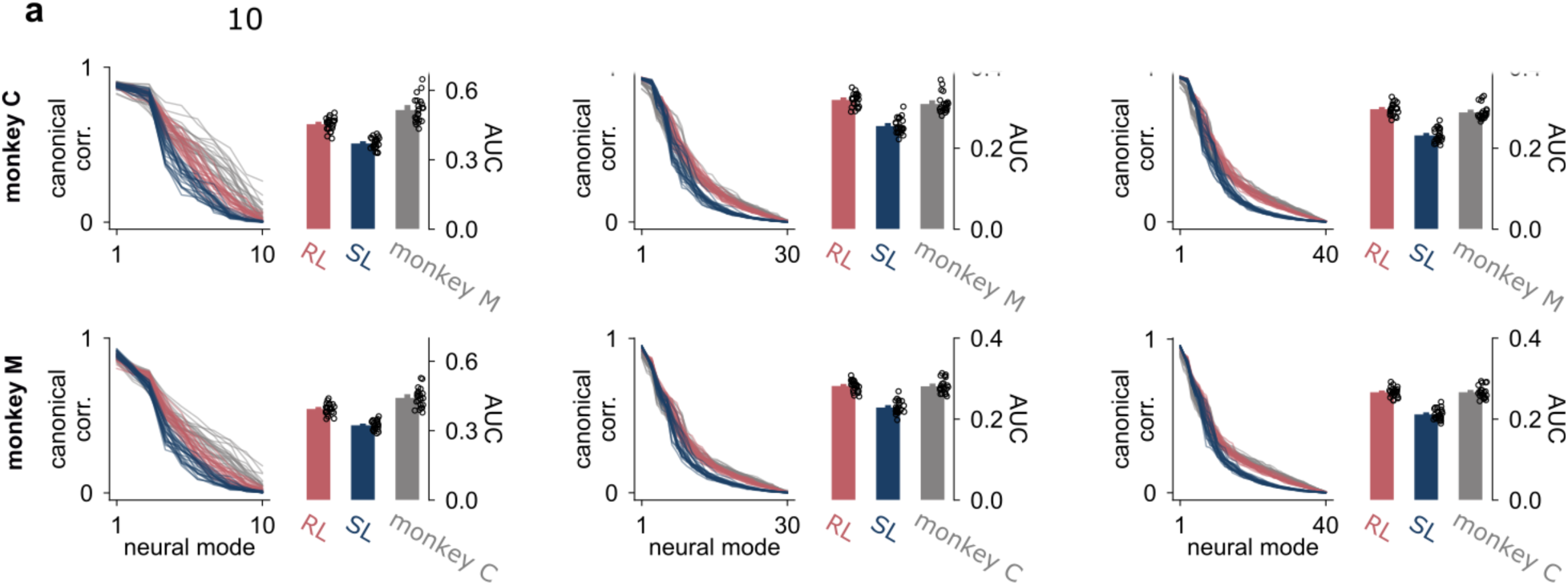
Varying the number of Principal Components used to compute Canonical Correlation curves did not alter the main results. **a**. 10 Principal Components. **b**. 10 Principal Components. **c**. 10 Principal Components. For each panel, the left subpanel shows the CC curves of our 24 seed-matched networks against the same recording session for monkey C (top) and monkey M (bottom). The grey lines represent comparison to different recording sessions from the other monkey. The right subpanels show the area under curve (AUC) of the CC curves. Bar heights and error bars indicate the mean and 95% confidence intervals, respectively.

**Fig. S7:**
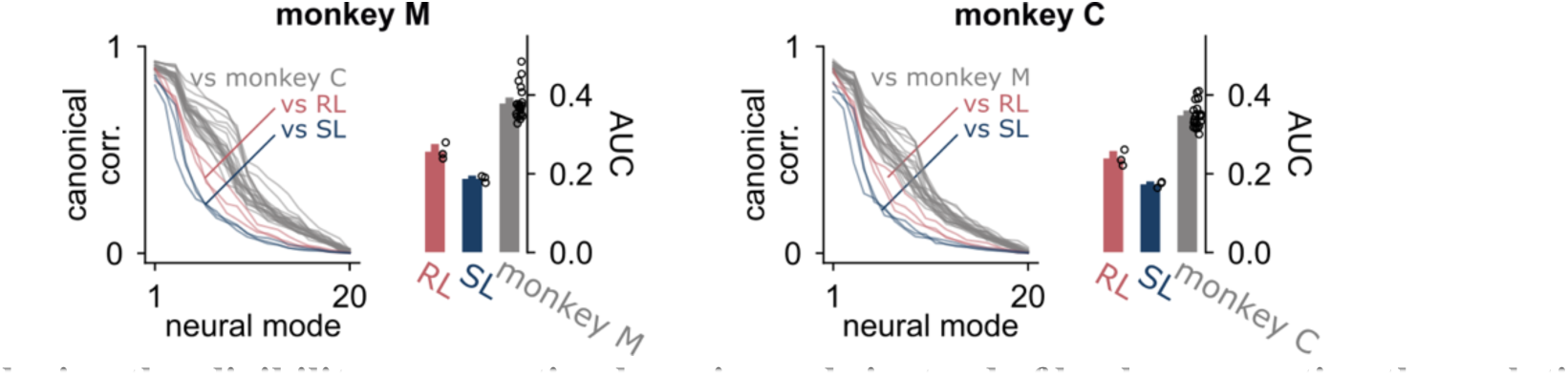
Employing the eligibility propagation learning rule instead of backpropagation through time did not alter the main results. For each panel, the left subpanel shows the CC curves of three seed-matched networks against the same recording session for monkey C (left) and monkey M (right). The grey lines represent comparison to different recording sessions from the other monkey. The right subpanels show the area under curve (AUC) of the CC curves. Bar heights and error bars indicate the mean and 95% confidence intervals, respectively.

**Fig. S8:**
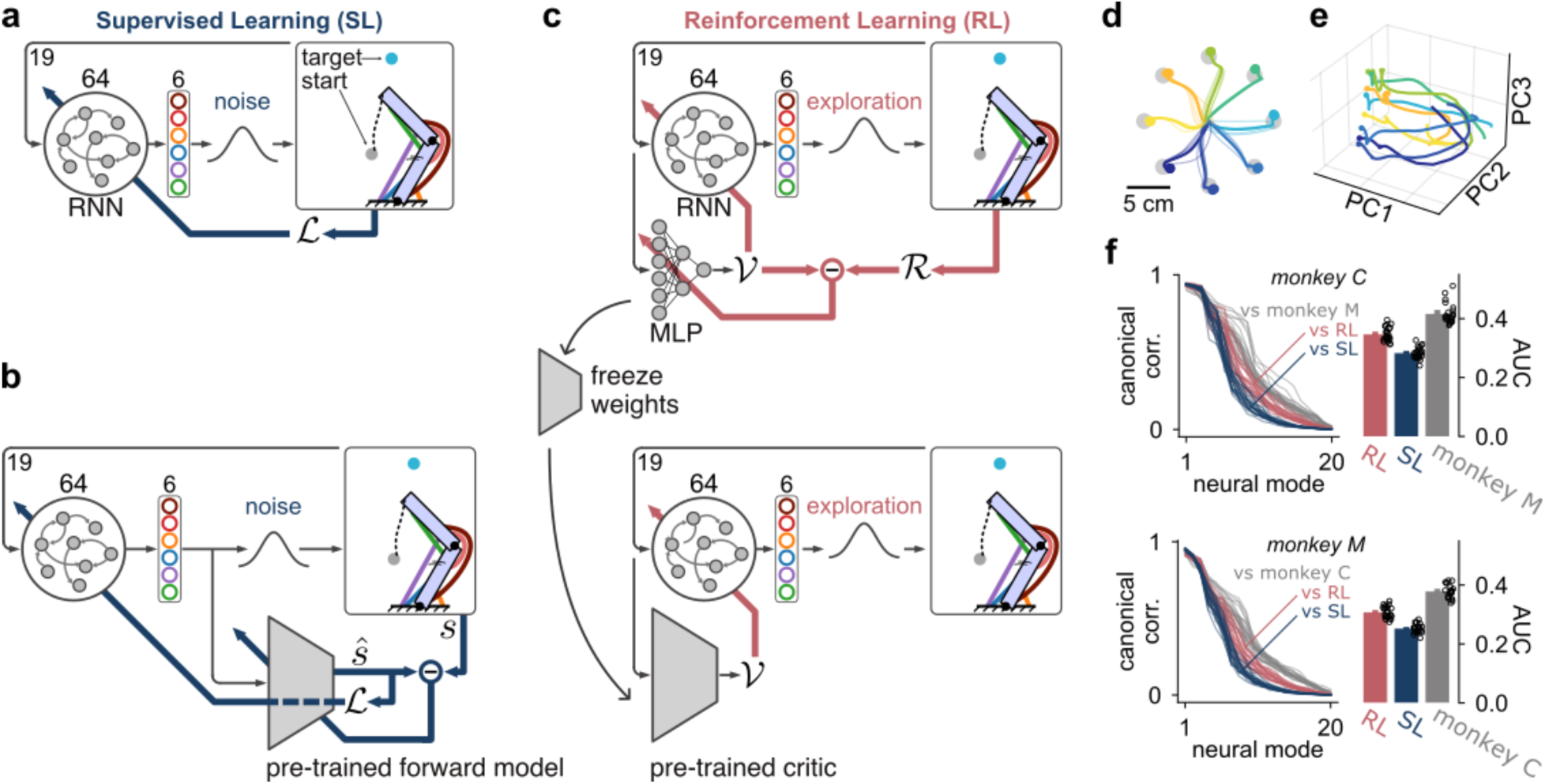
Pre-training the critic network does not alter the main results. **a.** The supervised learning architecture employed is conceptually equivalent to **b.**, a supervised learning architecture where a forward model predicts future effector states from the output actions driving the effector. Because the forward model is fully differentiable, the predicted states can be used to compute the loss and train the “actor” network with direct backpropagation through time. Therefore, the architecture in a. assumes a pre-trained forward model. **c.** In reinforcement learning, this can be mimicked with a pre-trained critic. We took the critics from the original RL models, froze the weights and used that critic to train a new actor network initialized using the same seed as the original model from which the pre-trained critic was taken. **d.** Example center-out reaches from an RL model after training with a pre-trained critic. **e.** Top three PCs from an RL model after training with a pre-trained critic, colored by reach direction like in d. **f.** Canonical Correlation scores and area under curve (AUC) from the original SL models presented in Fig. 3 and the RL models trained with a pre-trained critic.

**Fig. S9:**
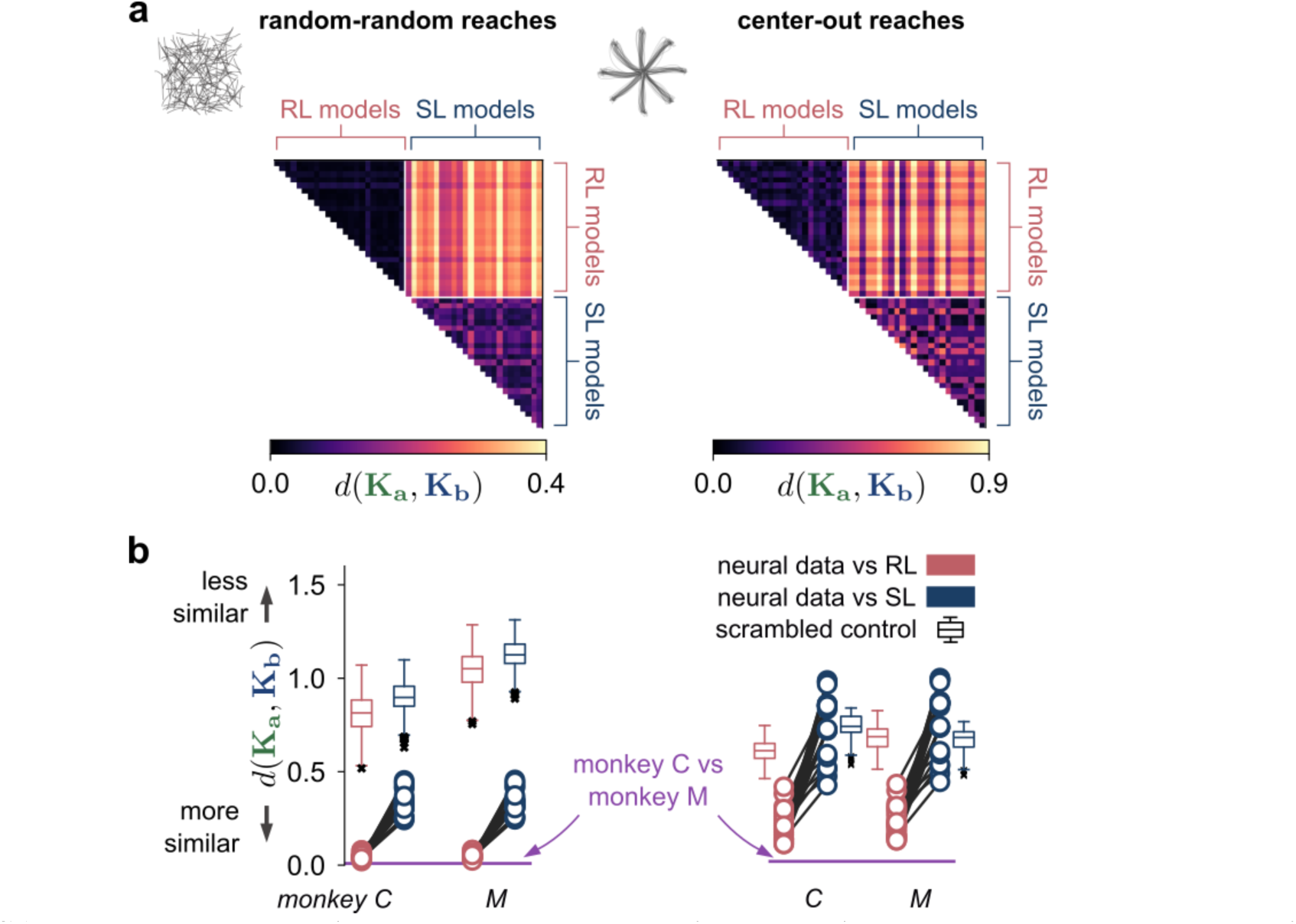
DSA on neural data during the center-out reaching task yielded the same results as during the random target task. **a**. DSA dissimilarity scores between neural networks models during a random target task (left subpanel, same data as in the main text) and center-out reaching task (right subpanel). Each row and column of the dissimilarity matrices represents a neural network seed. RL and SL seeds were ordered identically. **b**. Dissimilarity scores between neural network models and neural recordings from monkeys performing the random target task (left subpanel, same data as in the main text) and center-out reaching task (right subpanel). The solid horizontal purple line represents dissimilarity between the two monkey recordings.

**Fig. S10:**
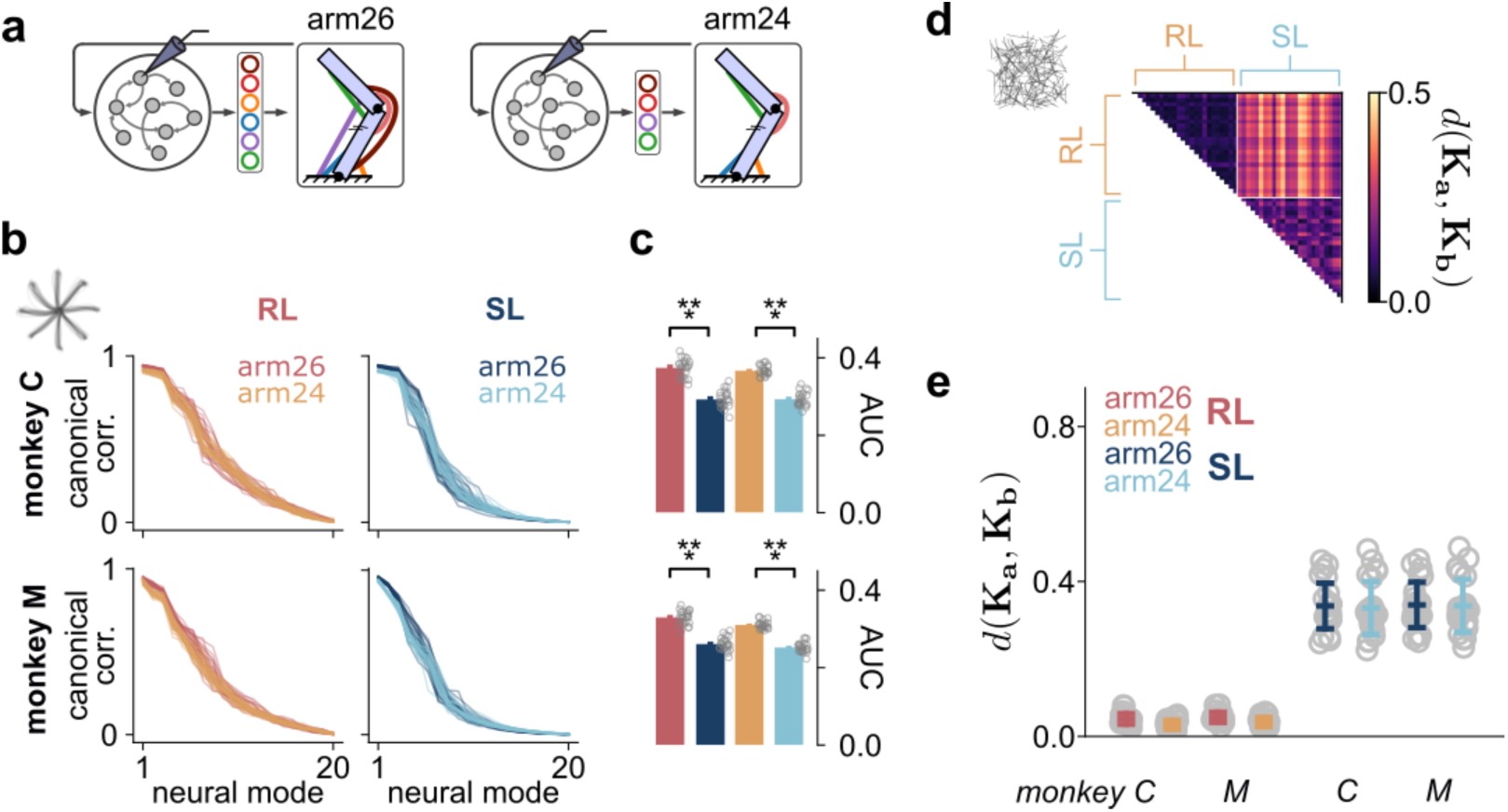
Dimensionality of the action vector did not alter the effect of RL training. **a**. We swapped the effector from a two-joints, six-actuators planar arm (arm26) with Hill-type muscles to a two-joints, four-actuators planar arm with Hill-type muscles (arm24). **b**, **c**. Learning to control an arm24 instead of an arm26 did not significantly alter Canonical Correlation scores between neural recordings and RL models during a center-out reaching task. AUC, area under curve. **d**. Dissimilarity scores were also significantly different between RL and SL models in a random target task. **e**. Dissimilarity scores against neural recordings indicate RL-trained neural networks still yield dynamical solutions much closer to neural recordings when trained to control an arm24 instead of an arm26. Bar height and error bars indicate mean and standard deviation, respectively.

**Fig S11:**
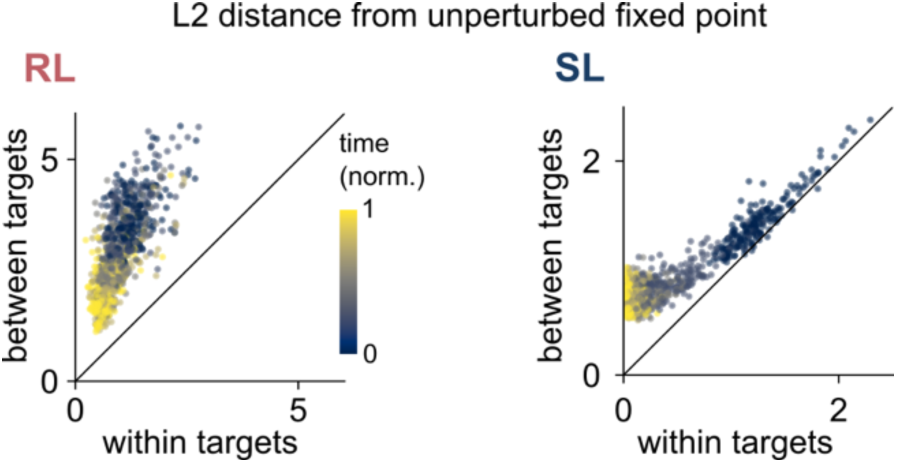
Neural distance from the unperturbed fixed point in a reach direction to all the perturbed fixed points from that same reach direction, plotted against unity against the neural distance to the perturbed fixed points in the other reaching directions. The points are color-coded based on when they occurred within a single reach, with 0 being the start of the reach and 1 being the end of the reach.

**Fig. S12:**
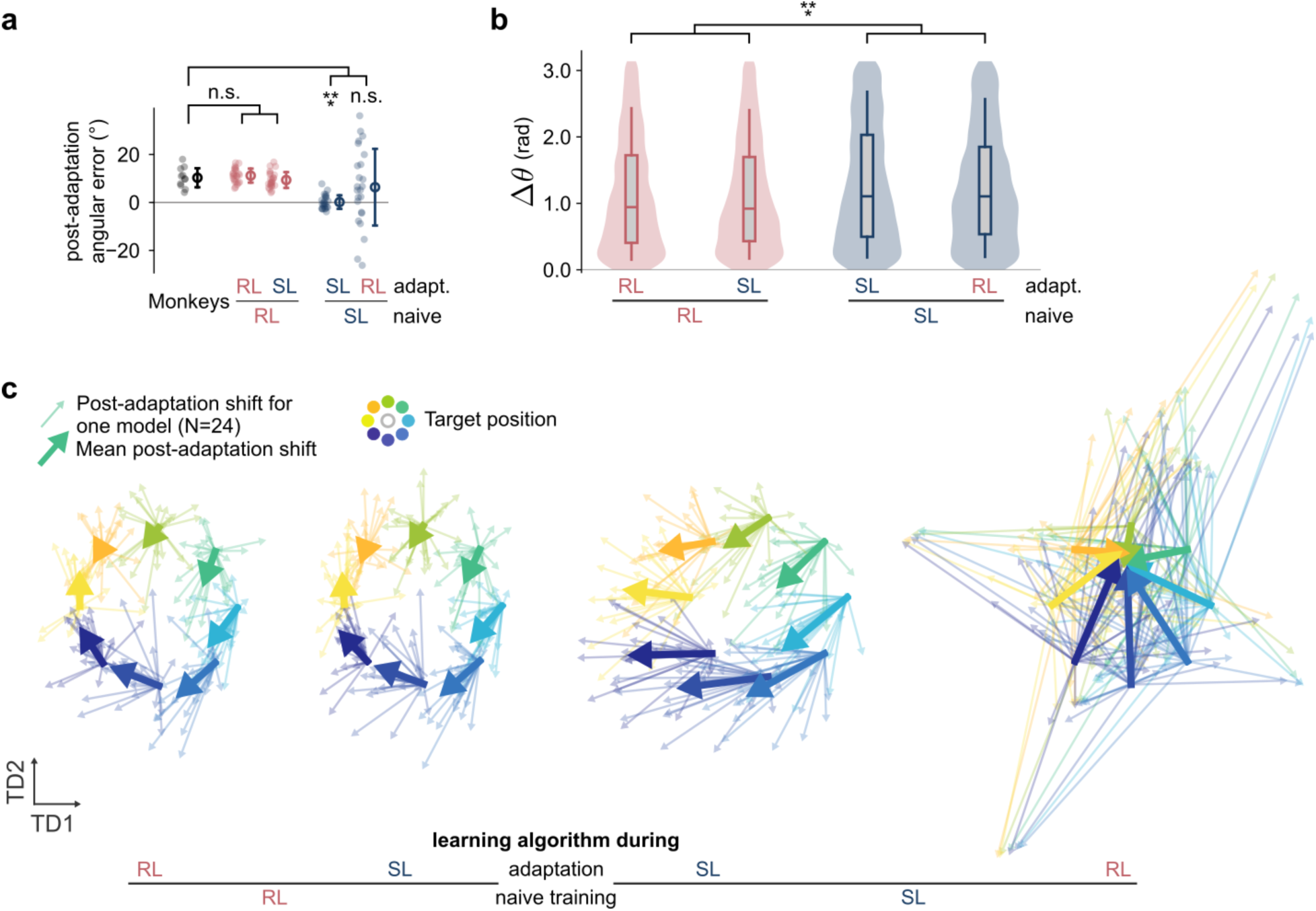
Neural reorganization induced by a visuomotor rotation after 300 training epochs. We analyzed neural network post-adaptation performance after 300 epochs instead of 500 epochs to assess if the results vary over adaptation. **a**. Angular deviation averaged over the last 8 trials of the adaptation block for each monkey session (black) and over thirty-two simulated trials after 300 epochs of adaptation for each neural network (24 seeds per group). The open circle and error bars indicate the mean and standard deviation, respectively. Angular error was quantified as mid-reach angular deviation from a straight trajectory, similar to the main text. Rank-sum test comparisons, RL-RL: U=0.53, p=0.59; RL-SL: U=0.85, p=0.39; SL-SL: U=4.55, p<0.001; SL-RL: U=1.07, p=0.29. **b**. Similarity between neural networks and neural recordings as the angular difference Δ*θ* between neural shifts in TDR subspace. Rank-sum test comparisons, RL-RL vs SL-SL: U=-5.12; RL-RL vs SL-RL: U=-4.58; RL-SL vs SL-SL: U=-5.40; RL-SL vs SL-RL: U=-4.96; p<0.001 for all comparisons. **c**. Shift of neural network activity in the target-predictive TDR subspace with arrows pointing from pre-to post-adaptation decoded states. Thick arrows represent across-session vector means. TDR: Targeted Dimensionality Reduction.

**Fig. S13:**
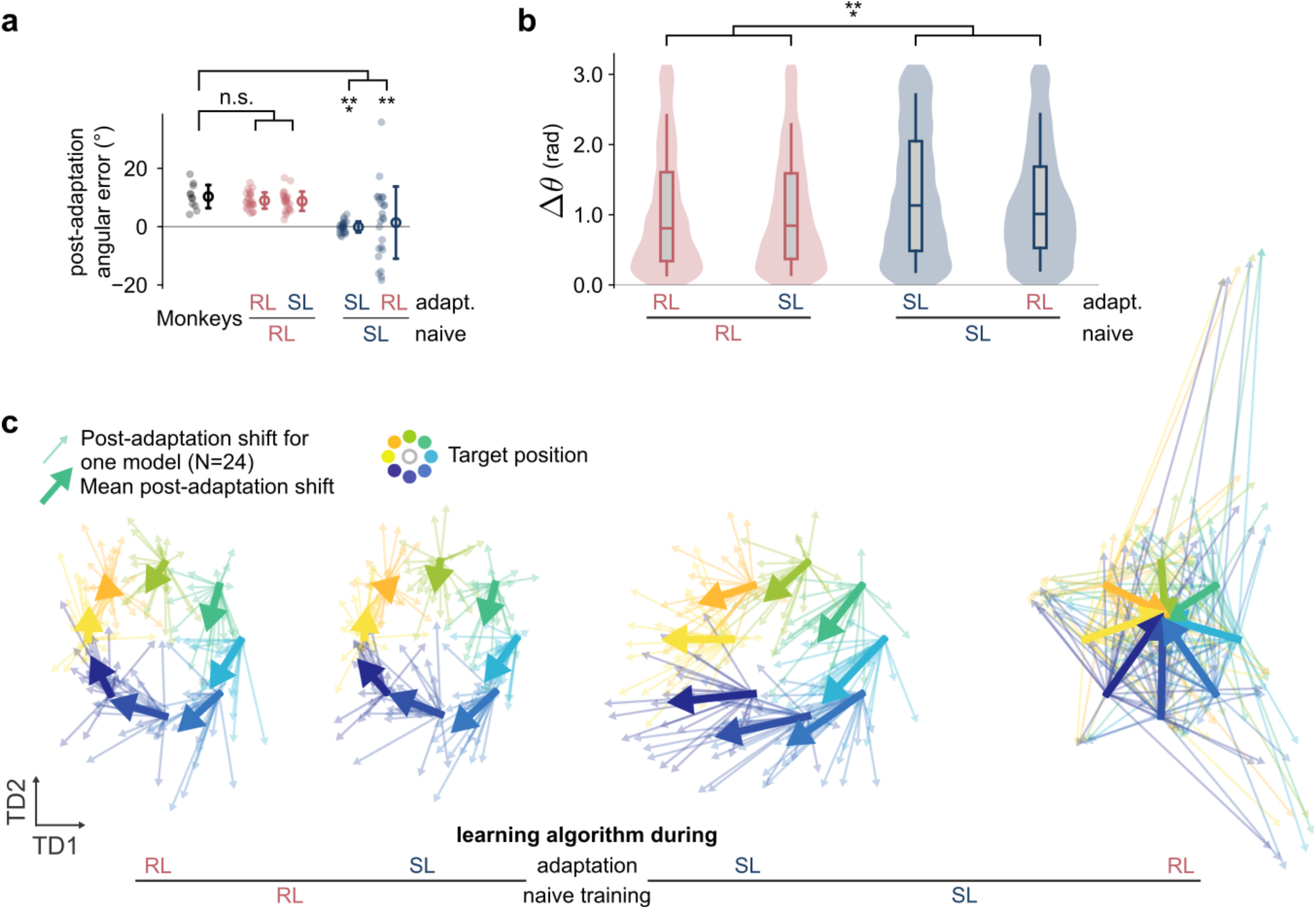
Neural reorganization induced by a visuomotor rotation with plastic input and recurrent weights. We trained our 24 seed-matched neural networks to adapt to a 30° visuomotor rotation while keeping both input and recurrent weights plastic, as opposed to just the input weights as in the main text. **a**. Angular deviation averaged over the last 8 trials of the adaptation block for each monkey session (black) and over thirty-two simulated trials after 500 epochs of adaptation for each neural network (24 seeds per group). The open circle and error bars indicate the mean and standard deviation, respectively. Angular error was quantified as mid-reach angular deviation from a straight trajectory, similar to the main text. Rank-sum test comparisons, RL-RL: U=1.10, p=0.27; RL-SL: U=0.89, p=0.37; SL-SL: U=4.65, p<0.001; SL-RL: U=2.59, p=0.009. **b**. Similarity between neural networks and neural recordings as the angular difference Δ*θ* between neural shifts in TDR subspace. Rank-sum test comparisons, RL-RL vs SL-SL: U=-8.77; RL-RL vs SL-RL: U=-6.24; RL-SL vs SL-SL: U=-8.38; RL-SL vs SL-RL: U=-5.55; p<0.001 for all comparisons. **c**. Shift of neural network activity in the target-predictive TDR subspace with arrows pointing from pre-to post-adaptation decoded states. Thick arrows represent across-session vector means. TDR: Targeted Dimensionality Reduction.

**Fig. S14:**
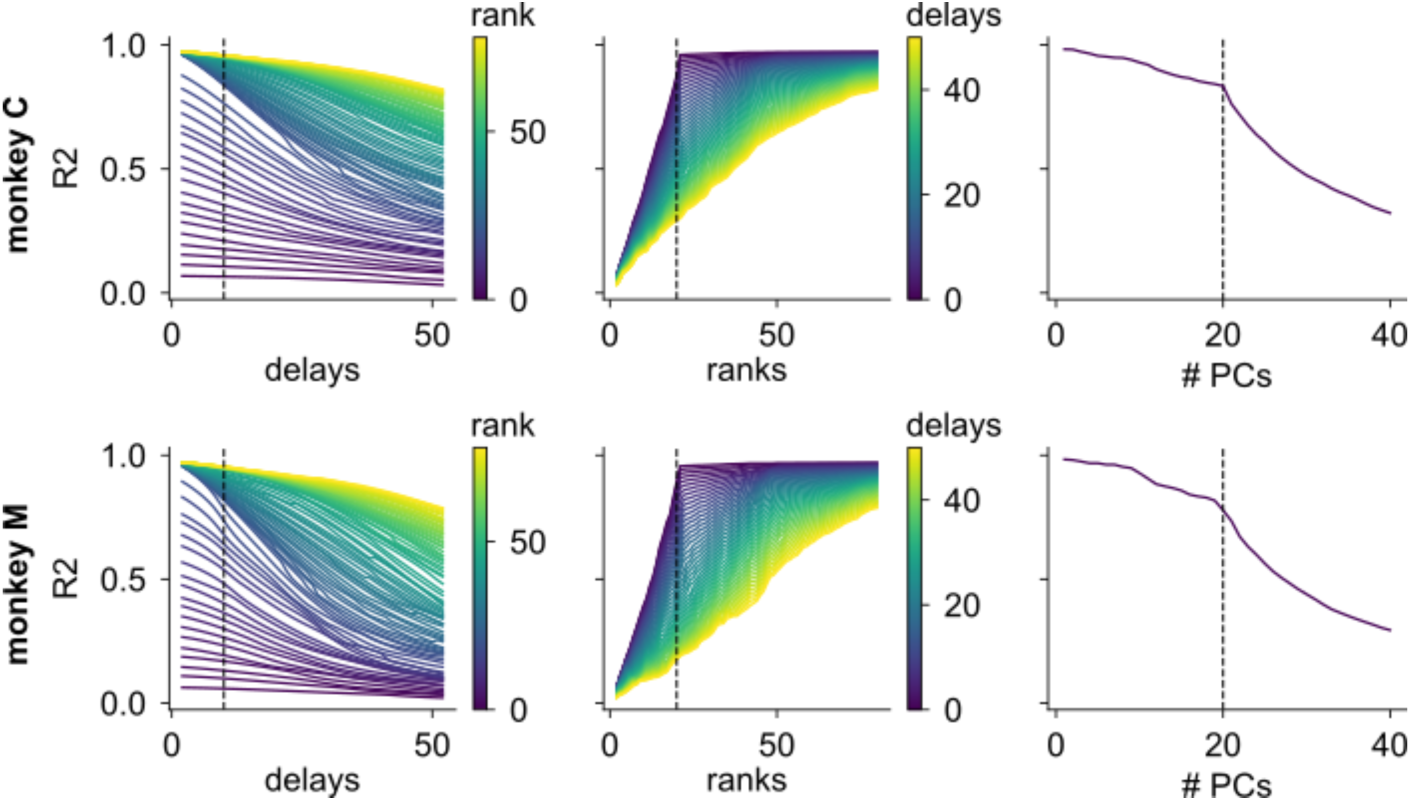
Dynamical Similarity Analysis hyperparameter sweep. Neural data for each monkey was divided into 10 folds following Principal Component dimensionality reduction, keeping the top 20 PCs similar to the Canonical Correlation analysis. The linearization map *g* was then computed over the first 9 folds, and the held-out data was projected into the resulting higher-dimensional subspace. The correlation R^2^ between the original data and its projection was then computed as we varied the hyperparameter values. Left column: sweep over number of delays for time-delayed embeddings. Middle column: sweep over number of regression ranks. We selected 10 delays and 20 ranks (see Methods for justification), indicated by the vertical dashed line on each plot. Right column: sweep over number of PCs for the selected hyperparameter

